# Benchmarking of alignment-free sequence comparison methods

**DOI:** 10.1101/611137

**Authors:** Andrzej Zielezinski, Hani Z. Girgis, Guillaume Bernard, Chris-Andre Leimeister, Kujin Tang, Thomas Dencker, Anna K. Lau, Sophie Röhling, JaeJin Choi, Michael S. Waterman, Matteo Comin, Sung-Hou Kim, Susana Vinga, Jonas S. Almeida, Cheong Xin Chan, Benjamin T. James, Fengzhu Sun, Burkhard Morgenstern, Wojciech M. Karlowski

**Affiliations:** Department of Computational Biology, Faculty of Biology, Adam Mickiewicz University in Poznan, Umultowska 89, 61-614 Poznan, Poland; Tandy School of Computer Science, The University of Tulsa, 800 South Tucker Drive, Tulsa, OK 74104, USA; Sorbonne Université, UMR 7205 ISYEB, Paris 75005 France; University of Göttingen, Institute of Microbiology and Genetics, Department of Bioinformatics, Goldschmidtstr. 1, 37077 Göttingen, Germany; Quantitative and Computational Biology Program, Department of Biological Sciences, University of Southern California, CA 90089, USA; Department of Chemistry, University of California, Berkeley, CA 94720; Molecular Biophysics & Integrated Bioimaging Division, Lawrence Berkeley National Laboratory, Berkeley, CA 94720; Department of Integrated Omics for Biomedical Sciences, Yonsei University, Seoul 03722, Republic of Korea; Korea Research Institute of Bioscience and Biotechnology, Daejeon 34141, Republic of Korea; Centre for Computational Systems Biology, School of Mathematical Sciences, Fudan University, Shanghai, 200433, China; Department of Information Engineering, University of Padova, Padova, Italy; INESC-ID, Instituto Superior Técnico, Universidade de Lisboa, Av. Rovisco Pais 1, 1049-001 Lisbon, Portugal; IDMEC, Instituto Superior Técnico, Universidade de Lisboa, Av. Rovisco Pais 1, 1049-001 Lisbon, Portugal; National Cancer Institute (NIH/NCI), Division of Epidemiology and Genetics (DCEG); Institute for Molecular Bioscience, and School of Chemistry and Molecular Biosciences, The University of Queensland, Brisbane, QLD 4072, Australia

**Keywords:** alignment-free, sequence comparison, benchmark, whole-genome phylogeny, horizontal gene transfer, web service

## Abstract

Alignment-free (AF) sequence comparison is attracting persistent interest driven by data-intensive applications. Hence, many AF procedures have been proposed in recent years, but a lack of a clearly defined benchmarking consensus hampers their performance assessment. Here, we present a community resource (http://afproject.org) to establish standards for comparing alignment-free approaches across different areas of sequence-based research. We characterize 74 AF methods available in 24 software tools for five research applications, namely, protein sequence classification, gene tree inference, regulatory element detection, genome-based phylogenetic inference and reconstruction of species trees under horizontal gene transfer and recombination events. The interactive web service allows researchers to explore the performance of alignment-free tools relevant to their data types and analytical goals. It also allows method developers to assess their own algorithms and compare them with current state-of-the-art tools, accelerating the development of new, more accurate AF solutions.

## BACKGROUND

Comparative analysis of DNA and amino acid sequences is of fundamental importance in biological research, particularly in molecular biology and genomics. It is the first and key step in molecular evolutionary analysis, gene function and regulatory region prediction, sequence assembly, homology searching, molecular structure prediction, gene discovery and protein structure-function relationships analysis. Traditionally, sequence comparison was based on pairwise or multiple sequence alignment (MSA). Software tools for sequence alignment, such as BLAST [1] and CLUSTAL [2], are the most widely used bioinformatics methods. Although alignment-based approaches generally remain the references for sequence comparison, MSA-based methods do not scale with the very large data sets that are available today. Additionally, alignment-based techniques have been shown to be inaccurate in scenarios of low sequence identity [3] (e.g., gene regulatory sequences [4,5] and distantly related protein homologs [3,6]). Moreover, alignment algorithms assume that the linear order of homologies is preserved within the compared sequences, so these algorithms cannot be directly applied in the presence of sequence rearrangements (e.g., recombination and protein domain swapping [7]) or horizontal transfer [8] in cases where large-scale sequence data sets are processed, e.g., for whole-genome phylogenetics [9]. In addition, aligning two long DNA sequences — millions of nucleotide long — is infeasible in practice. Therefore, as an alternative to sequence alignment, many so-called alignment-free (AF) approaches to sequence analysis have been developed [3], with the earliest works dating back to the mid 1970s [10], although the concept of the alignment-independent sequence comparison gained increased attention only in the beginning of the 2000s [11]. Most of these methods are based on word statistics or word comparison, and their scalability allows them to be applied to much larger data sets than conventional MSA-based methods.

A wide array of AF approaches to sequence comparison have been developed. These approaches include methods based on word or *k*-mer counts [12–16], the length of common substrings [17–20], micro-alignments [21–25], sequence representations based on chaos theory [26,27], moments of the positions of the nucleotides [28], Fourier transformations [29], information theory [30] and iterated-function systems [30,31]. Currently, the most widely used AF approaches are based on *k*-mer counts [32]. These methods are very diverse, providing a variety of statistical measures that are implemented across different software tools [3,33–35] (**Table 1**). Many *k*-mer methods work by projecting each of the input sequences into a feature space of *k*-mer counts, where sequence information is transformed into numerical values (e.g., *k*-mer frequencies) that can be used to calculate distances between all possible sequence pairs in a given data set.

**Table 1.** Alignment-free sequence comparison tools included in this study.

| Software | Approach class | Software version | Availability |
| --- | --- | --- | --- |
| AAF [72] | exact $k$-mer count | 10/01/2017 | https://github.com/fanhuan/AAF |
| AFKS [32] |  | 1.0 | https://github.com/TulsaBioinformaticsToolsmith/Alignment-Free-Kmer-Statistics |
| alfpy [3] |  | 1.0.6 | https://github.com/aziele/alfpy |
| CAFÉ [34] |  | 1.0.0 | https://github.com/younglululu/CAFE |
| FFP [33,38] |  | 2v.2.1 | https://github.com/jaejinchoi/FFP |
| jD2Stat [35] |  | 1.0 | http://bioinformatics.org.au/tools/jD2Stat/ |
| LZW-Kernel [79] | information theory | NA | https://github.com/kfattila/LZW-Kernel |
| spaced [84–86] | inexact $k$-mer count | 1.0 | http://spaced.gobics.de |
| kWIP [78] | $k$-mer count | 0.2.0-13-g3cf8a9e | https://github.com/kdmurray91/kWIP |
| ALFRED-G [57] | maximal length of exact common substrings | NA | https://alurulab.cc.gatech.edu/phylo |
| kmacs [18,85] |  | 1.0 | http://kmacs.gobics.de |
| kr [49] |  | 2.0.2 | http://guanine.evolbio.mpg.de/cgi-bin/kr2/kr.cgi.pl |
| Underlying Approach [90] |  | NA | http://www.dei.unipd.it/~ciompin/main/underlying.html |
| andi [22] | micro-alignments | 0.02 | https://github.com/EvolBioInf/andi |
| co-phylog [21] |  | NA | https://github.com/yhg926/co-phylog |
| FSWM [24]/Read-SpaM [76] |  | 1.0 | http://fswm.gobics.de |
| Multi-SpaM [23] |  | 1.0 | https://github.com/tdencker/multi-SpaM |
| phylonium |  | 0.3 | https://github.com/kloetzl/phylonium |
| mash [9] |  | 2.1 | https://github.com/marbl/Mash |
| Slope-SpaM | number of word matches | 0.1 | https://github.com/burkhard-morgenstern/Slope-SpaM |
| Skmer [52] |  | 3.0.0 | https://github.com/shahab-sarmashghi/Skmer |
| RTD-Phylogeny [82] | return time distribution | 1.0.1 | https://github.com/pandurang-kolekar/rtd-phylogeny |
| kSNP3 [77] | SNP count | 3.1 | https://sourceforge.net/projects/ksnp/files/ |
| EP-sim [73] | variable-length word counts | 1.0 | http://www.dei.unipd.it/~ciompin/main/EP-sim.html |

Detailed information about the tools parameter values used in this study for different reference data sets is provided in Additional file: **Table 1**. A concise description of the listed tools is provided in the **Methods**.

Despite the extensive progress achieved in the field of AF sequence comparison [3], developers and users of AF methods face several difficulties. New AF methods are usually evaluated by their authors, and the results are published together with these new methods. Therefore, it is difficult to compare the performance of these tools since they are based on inconsistent evaluation strategies, varying benchmarking data sets and variable testing criteria. Moreover, new methods are usually evaluated with relatively small data sets selected by their authors, and they are compared with a very limited set of alternative AF approaches. As a consequence, the assessment of new algorithms by individual researchers presently consumes a substantial amount of time and computational resources, compounded by the unintended biases of partial comparison. To date, no comprehensive benchmarking platform has been established for AF sequence comparison to select algorithms for different sequence types (e.g., genes, proteins, regulatory elements, or genomes) under different evolutionary scenarios (e.g., high mutability or horizontal gene transfer (HGT)). As a result, users of these methods cannot easily identify appropriate tools for the problems at hand and are instead often confused by a plethora of existing programs of unclear applicability to their study. Finally, as for other software tools in bioinformatics, the results of most AF tools strongly depend on the specified parameter values. For many AF methods, the word length *k* is a crucial parameter. However, words are used in different ways by different AF methods. Therefore, there is no universal word length *k* for all AF programs. Instead, different optimal word lengths have to be identified for the different methods. In addition, best parameter values may depend on the data-analysis task at hand, for instance, whether a set of protein sequences is to be grouped into protein families or superfamilies.

To address these problems, we developed AFproject (http://afproject.org), a publicly available web-based service for comprehensive and unbiased evaluation of AF tools. The service is based on eight well-established and widely used reference sequence data sets as well as four new data sets. It can be used to comprehensively evaluate AF methods under five different sequence analysis scenarios: protein sequence classification, gene tree inference, regulatory sequence identification, genome-based phylogenetics and HGT (**Table 2**). To evaluate the existing AF methods with these data sets, we asked the developers of 24 AF tools to run their software on our data sets or to recommend suitable input parameter values appropriate for each data set. In total, our study involved 10,202 program runs, resulting in 1,020,493,359 pairwise sequence comparisons (**Table 1**; Additional file 1: **Table S1**). All benchmarking results are stored and can be downloaded, reproduced and inspected with the AFproject website. Thus, any future evaluation results can be seamlessly compared to the existing ones obtained using the same reference data sets with precisely defined software parameters. By providing a way to automatically include new methods and to disseminate their results publicly, we aim to maintain an up-to-date and comprehensive assessment of state-of-the-art AF tools, allowing contributions and continuous updates by all developers of AF-based methods.

**Table 2.** Overview of the reference data sets.

| Category | Name | # Sequences | Average sequence length | # Files | # Sequence comparisons |
| --- | --- | --- | --- | --- | --- |
| Regulatory elements detection | Cis-regulatory modules (CRMs) [4] | 370 | 764 nt | 370 | 68,256 |
| Protein sequence classification | Low sequence identity (<40%) [93] | 1,066 | 180 aa | 1,066 | 567,645 |
| | High sequence identity (≥40%) [93] | 2,128 | 184 aa | 2,128 | 2,263,128 |
| Gene trees Inference | SwissTree [43] | 651 | 398 aa | 651 | 211,575 |
| Genome-based phylogeny | *Assembled genomes:* |  |  |  |  |
|  | 29 *E. coli*/Shigella strains | 29 | 4,895,247 nt | 29 | 406 |
|  | 14 plant species | 14 | 337,515,688 nt | 14 | 91 |
|  | 25 fish mitochondrial genomes [48] | 25 | 16,623 nt | 25 | 300 |
|  | *Unassembled genomes* |  |  |  |  |
|  | 29 *E. coli*/Shigella strains |  |  |  |  |
|  | coverage 0.03125 | 29,557 | 150 nt | 29 | 406 |
|  | coverage 0.0625 | 59,116 | 150 nt | 29 | 406 |
|  | coverage 0.125 | 118,266 | 150 nt | 29 | 406 |
|  | coverage 0.25 | 236,541 | 150 nt | 29 | 406 |
|  | coverage 0.5 | 473,081 | 150 nt | 29 | 406 |
|  | coverage 1 | 946,169 | 150 nt | 29 | 406 |
|  | coverage 5 | 4,730,778 | 150 nt | 29 | 406 |
|  | 14 plant species |  |  |  |  |
|  | coverage 0.015625 | 48,274 | 150 nt | 14 | 91 |
|  | coverage 0.03125 | 96,489 | 150 nt | 14 | 91 |
|  | coverage 0.0625 | 1,931,268 | 150 nt | 14 | 91 |
|  | coverage 0.125 | 3,862,905 | 150 nt | 14 | 91 |
|  | coverage 0.25 | 7,725,928 | 150 nt | 14 | 91 |
|  | coverage 0.5 | 15,461,718 | 150 nt | 14 | 91 |
|  | coverage 1 | 30,903,727 | 150 nt | 14 | 91 |
| Horizontal gene transfer | 27 *E. coli/Shigella* genomes [54] | 27 | 4,905,896 nt | 27 | 351 |
|  | 8 *Yersinia* species [56] | 8 | 4,605,553 nt | 8 | 28 |
|  | 33 simulated genomes [53] |  |  |  |  |
|  | HGT level 0 | 33 | 2,205,524 nt | 33 | 528 |
|  | HGT level 250 | 33 | 2,149,620 nt | 33 | 528 |
|  | HGT level 500 | 33 | 2,230,317 nt | 33 | 528 |
|  | HGT level 750 | 33 | 2,263,926 nt | 33 | 528 |
|  | HGT level 1,000 | 33 | 2,238,661 nt | 33 | 528 |

An interactive visualization of all results for all data sets can be found online (http://afproject.org).

## RESULTS

### Benchmarking service

To automate AF method benchmarking with a wide range of reference data sets, we developed a publicly available web-based evaluation framework (**Figure 1**). Using this workflow, an AF method developer who wants to evaluate his/her own algorithm first downloads sequence data sets from one or more of the five categories (e.g., data set of protein sequences with low identity from the protein sequence classification category) from the server. The developer then uses the downloaded data set to calculate pairwise AF distances or dissimilarity scores between the sequences of the selected data sets. The benchmarking service accepts the resulting pairwise distances in tab-separated value (TSV) format or as a matrix of pairwise distances in standard PHYLIP format. In addition, benchmarking procedures in two categories (genome-based phylogeny and horizontal gene transfer) also support trees in Newick format to allow for further comparative analysis of tree topologies.

**Figure 1.**
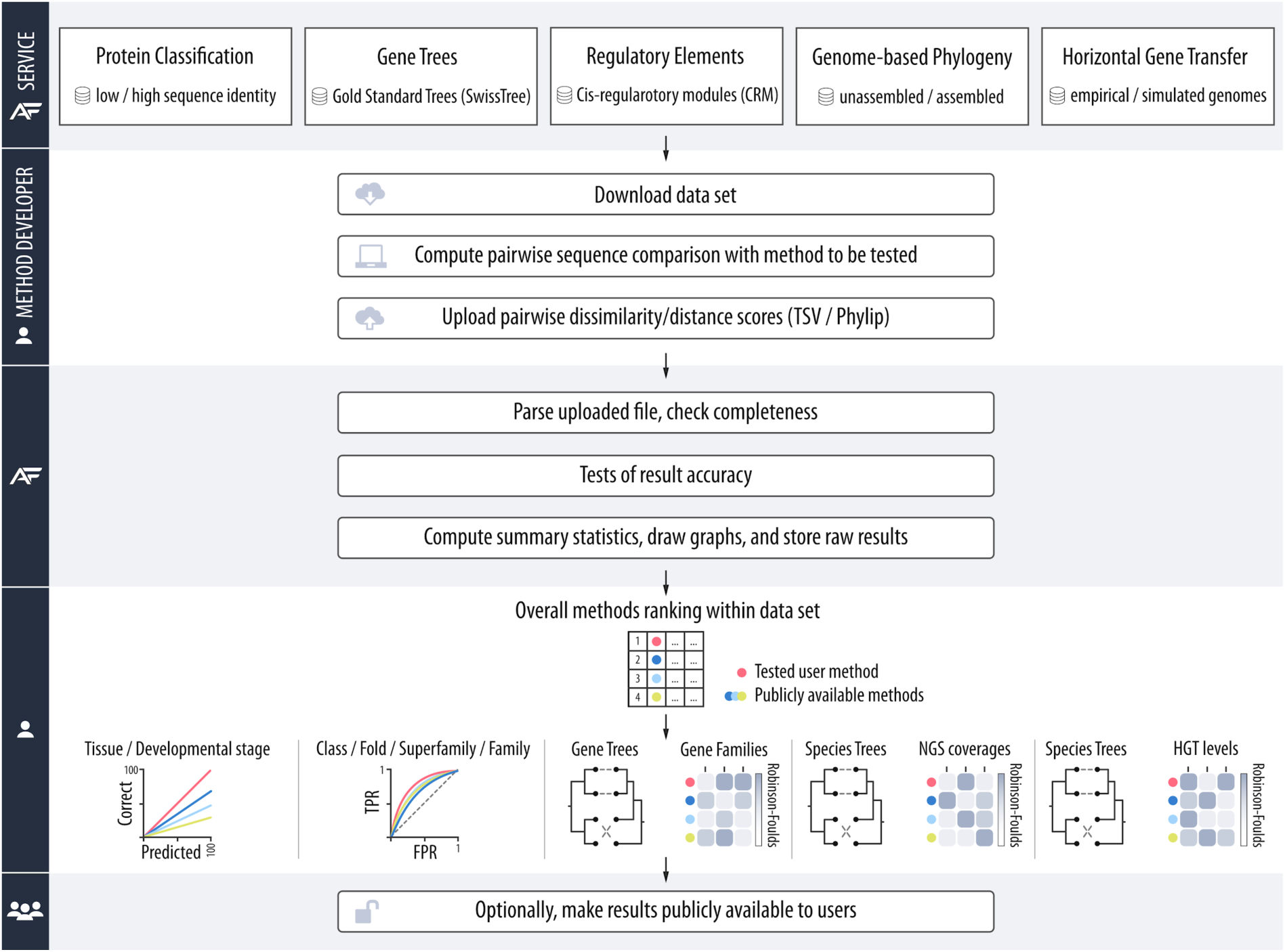
Overview of the AFproject benchmarking service facilitating assessment and comparison of AF methods. AF method developers run their methods on a reference sequence set and submit the computed pairwise sequence distances to the service. The submitted distances are subjected to a test specific to given data sets, and the results are returned to the method developer, who can choose to make the results publicly available.

Once the output file is uploaded to the AFproject web server, the service starts the benchmarking procedure, which is typically completed in a few seconds. Finally, the raw data and the time-stamped benchmark report are stored and provided to the submitter. The report shows the performance of the evaluated method and compares it with the performance of other methods that have been previously evaluated through the AFproject web server. In the report, the performance of the compared methods is ordered by a statistical measure specific to the respective benchmark category (e.g., the Robinson-Foulds distance measure [36] in the categories of gene trees, genome-based phylogeny and horizontal gene transfer). By default, the report is private (visible only to the submitter), and the developer can choose if and when to make the report publicly available. Similar to other benchmarking platforms [37], we have released the source code of the web service to facilitate transparency and encourage feedback and improvements from the community (https://github.com/afproject-org/afproject).

### Alignment-free method catalog

To evaluate the performance of currently available AF tools and create a reference data for future comparisons, we benchmarked 24 standalone tools (**Table 1**), covering a large proportion of the currently available AF methods. Some tools offer multiple related methods to measure pairwise distances (or dissimilarity) between sequences; for instance, jD2Stat [35] supports three different distance measures based on the D_2_ statistic: jD2Stat--d2n, jD2Stat--d2s and jD2Stat--d2st. In this study, we included these different distance measures, resulting in a total of 74 tested tool variants (**Figure 2**). Each of these tool variants was run with various combinations of parameter values (Additional file 1: **Table S1**). The values yielding the best performance for a given method were selected and saved in the AFproject database; if multiple parameters produced the same best-performing results for a tool, we selected only the values that were least computationally demanding (e.g., the shortest word length for word-counting methods or the smallest sketch size). Full information about the benchmarking results, including all combinations of parameter values of the evaluated tools, can be downloaded from http://afproject.org/download/.

**Figure 2.**
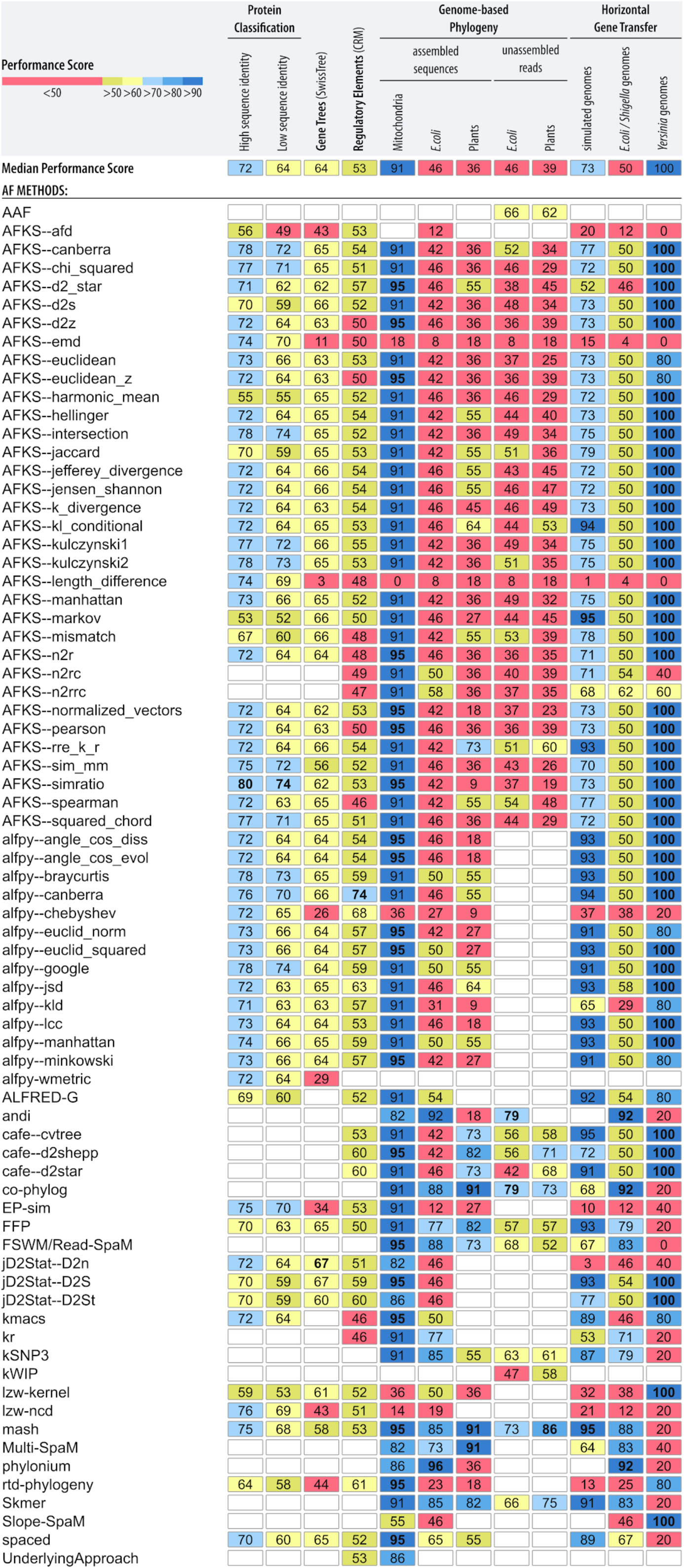
Summary of AF tool performance across all reference data sets. The numbers in the fields indicate the performance scores (from 0 to 100; see **Methods**) of a given AF method for a given data set. Fields are color-coded by performance values. The numbers in bold indicate the highest performance obtained within a given data set. An empty field indicates the corresponding tool’s inability to be run on a data set. An extended version of this figure including values of the overall performance score is provided in Additional file 1: **Table S14**. The most up-to-date summary of AF tool performance can be found at: http://afproject.org/app/tools/performance/.

Only three tools (Alignment-Free-Kmer-Statistics (AFKS) [32], FFP [38] and mash [9]) are sufficiently generic to be applied to all 12 benchmarking data sets; the remaining tools can handle only subsets of our reference data sets, either because they have been designed only for a specific purpose (e.g., to handle only certain sequence types, such as nucleotides, proteins, and unassembled or assembled genomic sequences) or — less frequently — because of some unexpected software behavior (e.g., a program stops functioning, does not terminate in a reasonable amount of time or produces invalid results; Additional file 1: **Table S1**). Hence, one of the results of our benchmarking study is an extensive and annotated catalog of tools (http://afproject.org/tools/), which constitutes a resource not only for users of AF methods, but also for the developers of these methods, as it should help identify which aspects of existing algorithms may be in need of further development.

### Protein sequence classification

Recognition of structural and evolutionary relationships among amino acid sequences is central to the understanding of the function and evolution of proteins. Historically, the first comprehensive evaluation of AF methods [6] investigated the accuracy of the tools for protein structure classification at four hierarchical levels used in the Structural Classification of Proteins (SCOP) database, namely, family, superfamily, class, and fold [39]. The original protocol tested six *k*-mer-based distance measures against a subset of the SCOP database, containing protein family members sharing less than 40% sequence identity [6]. In the present study, we extend the original analysis [6] to test the accuracy of 56 tool-measure variants in recognition of structural relationships of protein sequences sharing both low (<40%) and high (≥40%) sequence identity (**Figure 2**).

The area under the receiver operating characteristics (ROC) curve (AUC), which indicates whether a method is able to discriminate between homologous and nonhomologous protein sequences (**Methods**), showed the favorable performance of AFKS [32] software. AFKS with parameters set to the *simratio* [32] distance and a word length of *k* = 2 is the best performing tool for both low- and high-sequence-identity data sets (**Figure 2**). For the latter type of the data set, the method produces the highest AUC values across all four structural levels, with an average AUC of 0.798 ± 0.139 (Additional file 1: **Table S2**). When considering the low-sequence-identity data set (Additional file 1: **Table S3**), AFKS--*simratio* also has the highest average AUC of 0.742 ± 0.079 but lower performance at the superfamily and family levels than alfpy [3] (set to the Google distance and *k* = 1). alfpy--*google* is ranked *2*^nd^ (0.738 ± 0.091) and fourth (0.778 ± 0.142) for the low- and high-sequence-identity data sets, respectively. Notably, the top-seven-ranking positions in both the low- and high-sequence-identity data sets are occupied, though in a different order, by the same measures from AFKS and alfpy software (**Figure 2**).

In general, the tested tools achieve greater discriminatory power in recognizing structural relationships (higher average AUCs) in our high-sequence-identity data set than in the low-sequence-identity data set (**Figure 2**; Wilcoxon signed rank test: *p* = 2.602 × 10^−11^). Almost all tool variants, except AFKS--*afd* (AUC: 0.492 ± 0.016) for the low-sequence-identity data set, achieved higher overall performance than the random classifier (AUC > 0.5). As expected and previously reported [3,6], the tools lose discriminatory power from the family to the class level for both data sets (the AUC decreases; Additional file 1: **Tables S2-S3**), as the sequence similarity is lower within higher hierarchical groups. As a result, all methods tested (except AFKS--*harmonic_mean*) achieve their best accuracy at the family level. The AUC values at the family, superfamily and fold levels are higher (Wilcoxon signed rank tests: *p* < 10^−5^) for data sets with high sequence similarity than for data sets with low sequence similarity. The greatest difference in performance was observed at the family level, where the maximum AUC obtained by the tools with the high- and low-sequence-identity data sets was 1.0 and 0.84, respectively. The methods result in more similar AUCs at the class level for the low-sequence-identity data set than for the high-sequence-identity data set (Wilcoxon signed rank tests: *p* = 0.0185). Protein sequences at the class level lack conserved segments, and the median AUC values obtained by the methods in high- and low-sequence-identity data sets are similar to those obtained with the random classifier (median AUC: 0.57 in both data sets).

### Gene tree inference

Only a few studies [40,41] have evaluated AF methods in the construction of gene trees. Because of the limited amount of sequence information available, gene trees are typically more difficult to reconstruct than species trees [42]. We assessed the accuracy of 11 AF tools (55 tool variants) in inferring phylogenetic relationships of homologous sequences based on a collection of high-confidence SwissTree phylogenies representing different types of challenges for homology prediction, e.g., numerous gene duplications and HGT [37,43]. Similar to SwissTree, we assessed the gene families at the protein-sequence level to minimize the impact of codon degeneracy. We thus interpret an inferred phylogenetic tree based on a homologous family of protein sequences as the tree for the gene family (i.e., the gene tree). As a measure of accuracy, we computed the normalized Robinson-Foulds (nRF) distance [36] between the trees reconstructed by the AF methods under study and the reference trees. The nRF distance has values between 0 and 1, with 0 indicating identical tree topologies and 1 indicating the most dissimilar topologies.

None of the AF methods that we tested were able to perfectly infer the respective reference tree topology for any of the 11 gene families. jD2Stat [35] (D_2_^n^ with parameter values *n*=1 and *k*=5) was the most accurate tool in our test (**Figure 2**). This method achieved the lowest nRF values (highest accuracy) among all the tested methods averaged across all 11 reference gene families (nRF = 0.3296 ± 0.1511; Additional file 1: **Table S4**), which can be interpreted as 33% (± 15%) of incongruent bipartitions between the inferred and the reference tree. To put this number into perspective, the corresponding gene trees based on MSA (i.e., neighbor-joining trees inferred using ClustalW alignments generated with default parameters) yielded a similar average accuracy (nRF = 0.2995 ± 0.1511). In general, the nRF distances obtained by the tested methods vary greatly across the gene families (Friedman rank sum test: *p* < 2.2 × 10^−16^, *df* = 10, Friedman chi-square = 463.88) due to different complexities of the encoded protein families (e.g., evolutionary distance between proteins, domain architecture, and structural and functional affiliations). Consequently, the tools obtain their best accuracy in phylogenetic inference of the eukaryotic protein family of sulfatase modifying factor (SUMF) proteins, which are characterized by a single protein domain and the smallest number of gene duplications; four distance measures in AFKS software generated trees (nRF = 0.077) with minor topological differences in the speciation order of three proteins (Additional file 2: **Figure S1**). The AF methods achieved the second-best accuracy (median nRF = 0.178) for the eukaryotic NOX family NADPH oxidases — a gene family coding for transmembrane enzymes with 10 gene duplications and 3–4 protein domains. However, the examined tools produced highly inaccurate phylogenetic trees of two other transmembrane protein families, namely, Bambi and Asterix (median nRFs: 0.615 and 0.611, respectively), where more than 60% of tree topologies differed from the reference tree.

### Regulatory elements

Analysis of gene regulatory sequences is another domain where AF methods are popular, as the similarity between these elements is usually low and alignments typically fail to detect it properly [4]. We adopted a benchmarking procedure and a reference data set of cis-regulatory modules (CRMs) introduced by Kantarovitz et al. [4], which was further used in other studies [44], showing that alignment algorithms lag behind AF methods in recognizing functionally-related CRMs. A CRM can be broadly defined as a contiguous noncoding sequence that contains multiple transcription factor binding sites and regulates the expression of a gene. The Kantorovitz’ protocol assesses to what extent AF tools are capable of capturing the similarities between functionally-related CRMs, expressed in the tissues of fly and human (see **Methods**).

However, none of the AF methods produced perfect results for any of the seven tissues/species data set combinations (i.e., all functionally related CRM pairs classified in front of all random DNA pairs). alfpy software [3] set to three distance measures — Canberra, Chebyshev and Jensen-Shannon divergence — captured the largest number (averaged across 7 tissue samples) of functionally related regulatory elements (**Figure 2**). The selection of Canberra distance (word length of *k* = 2) correctly recognized 73.6% ± 10.54% of CRMs, capturing the highest functional relatedness in three out of the seven data sets (tracheal system: 97%, eye: 78% and blastoderm-stage embryo: 76% in fly; Additional file 1: **Table S5**). The Chebyshev distance (*k* = 7) obtained the second-highest average performance of 67.59% and the highest performance variation across seven data sets (standard deviation = 20.14%) among all methods in the ranking; this measure had the highest performance for two tissues (peripheral nervous system in fly and HBB complex in human) and relatively low performance in human liver tissue. The third measure, Jensen-Shannon divergence (*k* = 2), achieved more stable performance across the data sets than the Canberra and Chebyshev distances (63.16% ± 8.22%). Overall, 51 out of 63 methods showed average performance better than that of the random classifier (> 50%).

### Genome-based phylogeny

AF methods are particularly popular in genome-based phylogenetic studies [9,12,13,38] because of (i) the considerable size of the input data and (ii) complex correspondence of the sequence parts, often resulting from genome rearrangements [45]. Additionally, no statistical substitution models are currently available for assessing the evolution of complete genomes. We assessed the ability of AF methods to infer species trees using benchmarking data from different taxonomic groups, including bacteria, animals and plants. Here, we used completely assembled genomes as well as simulated unassembled next-generation sequencing reads at different levels of coverage.

#### Assembled genomes

As many studies have applied AF methods to whole mitochondrial genomes [46,47], we tested the performance of 23 AF software tools (70 tool variants in total) in phylogenetic inference using complete mtDNA from 25 fish species of the suborder Labroidei [48]. The best accuracy was achieved by nine AF tools (19 tool variants), which generated tree topologies that were almost identical to the reference Labroidei tree (nRF = 0.05; **Figure 2**; Additional file 1: **Table S6**). The results differ only in the speciation order of three closely related fish species belonging to the Tropheini tribe of the Pseudocrenilabrinae family (Additional file 2: **Figure S2**). The same species were misplaced in the topologies generated by additional 39 tool variants that all occupied the second place in the benchmark ranking (nRF = 0.09). These methods additionally misplace species within the Pomacentridae and Embiotocidae families. These results indicate that most AF methods infer trees in general agreement with the reference tree of mitochondrial genomes [18,46,49,50].

We further tested the performance of AF methods in phylogenetic inference with larger, bacterial genomes of *Escherichia coli/Shigella* and with nuclear genomes of plant species (**Figure 2**). Seven tools (nine tool variants) could not be tested on all three sets of complete genomes since the programs did not complete analyses (Additional file 1: **Table S1**). The remaining 16 tools (61 tool variants) led to greater nRF distances, i.e., lower performance for the phylogeny of the *E. coli/Shigella* and plant nuclear genomes than for the phylogeny of mitochondrial genomes (**Figure 2**; one-way analysis of variance (ANOVA) with repeated measures: *p* < 2 × 10^−16^; *post hoc* pairwise paired t test: *p* < 2× 10^−16^). Although the tools that we tested show similar nRF distances for bacterial and plant genomes in general (pairwise paired t test: *p* = 0.073), the top-performing tools are different between the two data sets. For example, phylonium [51] and andi [22], which were developed for phylogenetic comparison of closely related organisms, are the best performing tools for the *E. coli/Shigella* data sets, whereas on the plant data sets, both tools perform poorly (**Figure 2**). Phylonium almost perfectly reproduced the reference tree for the *E. coli/Shigella* group with an nRF = 0.04 (Additional file 1: **Table S7**; there was only a single error in the placement of two closely related *E. coli* K-12 substrains: BW2952 and DH10B; Additional file 2: **Figure S3**), while the plant trees obtained by these tools showed very low topological similarity to the reference tree (nRF = 0.64; Additional file 1: **Table S8**).

The best-performing tools for the plant data set are co-phylog [21], mash [9] and Multi-SpaM [23], all of which almost perfectly recovered the reference tree topology of the plant species (with an nRF = 0.09 for all three programs). In each of the trees produced by these programs, there was exactly one species placed at an incorrect position compared to its position in the reference tree, namely, in the speciation order in the Brassicaceae family for co-phylog (Additional file 2: **Figure S4**), mash (Additional file 2: **Figure S5**) and for Multi-SpaM, the last of which placed *Carica papaya* outside the Brassicales order (Additional file 2: **Figure S6**). Additionally, co-phylog is the third-best-performing tool in reconstructing the *E. coli/Shigella* tree topology (nRF = 0.12), while mash and Multi-SpaM are at the fourth and sixth positions, respectively, in this ranking (nRF = 0.15 and nRF = 0.27, respectively). As a result, co-phylog, mash, FFP [33], Skmer [52] and FSWM [24] are among the top 5 best-performing tools for both data sets (**Figure 2**),.

#### Raw sequencing reads

We also tested the accuracy of AF tools in phylogenetic inference based on unassembled sequencing reads, represented by seven different levels of sequencing coverage, from *E. coli/Shigella* and from a set of plant species (**Table 1**; see **Methods**). No differences in nRF values were observed between the results based on the unassembled and assembled *E. coli/Shigella* genomes (Wilcoxon signed rank test: *p* = 0.169), indicating that the AF tools exhibited equal performance for the unassembled and assembled genomes. In contrast, the tested tools showed lower performance (i.e., higher nRF values) in assembly-free phylogenetic reconstruction of the plant species (Wilcoxon signed rank test: *p* = 0.00026). andi and co-phylog [21] are the most accurate tools in the *E. coli/Shigella* data set (**Figure 2**), with an average nRF distance of 0.21 ± 0.14 (Additional file 1: **Table S9**). Both tools achieved the minimum nRF for seven coverage levels in the *E. coli/Shigella* data set (i.e., andi for coverage 0.03125, 0.25, 0.5 and 5, and co-phylog for coverage from 0.0625 to 0.125 and from 1 to 5). Although andi could not be tested with unassembled plant data set due to high sequence divergence (Additional file: **Table S1**), the accuracy of co-phylog for this set is similar as for *E. coli/Shigella* data (nRF = 0.27 ± 0.13; Additional file 1: **Table S10**), which places the tool at the 3^rd^ position in the ranking for the plant sequences (**Figure 2**).

For the unassembled plant data sets, mash is the most accurate tool (**Figure 2**), i.e., the tool with the shortest nRF distance between the inferred trees and the reference tree. For the lowest coverage level (0.015625), mash still allows us to infer trees with average nRF distances of 0.27 from the reference tree (Additional file 1: **Table S10**). In general, mash shows best performance at six out of the seven coverage levels (i.e., from 0.015625 to 0.5). For the unassembled *E. coli/Shigella* data set, mash is ranked at the 2^nd^ position, with an average nRF distance of 0.27 ± 0.18. Notably, for coverage 0.25 in plant data set, mash inferred tree topology in perfect agreement with the reference tree (nRF = 0; Additional file 1: **Table S10**), however its performance slightly decreases for higher coverage levels (with nRFs of 0.09 and 0.18 for coverage 0.5 and 1, respectively). The best accuracy at the highest coverage level (1x) was obtained by co-phylog (nRF = 0.09).

When considering the most universal tools applied to all the tested reference data sets, mash ranks first and the second for the assembly-free phylogeny of plants and *E. coli/Shigella*, respectively (**Figure 2**). In addition to mash, two other methods designed specifically for phylogenetic reconstruction from next-generation sequencing data — co-phylog and Skmr — are the only tools ranked among the top 5 methods tested on both unassembled data sets (**Figure 2**).

### Horizontal gene transfer

To assess the accuracy of the AF methods in phylogenetic reconstruction of sequences that underwent frequent HGT events and genome rearrangements, we used sets of simulated genomes with different levels of HGT [53] as well as two real-world data sets of microbial species, namely, 27 genomes of *E. coli* and *Shigella* [53–55] and eight *Yersinia* genomes [53,56] (**Table 1**). Similarly to previous tests, we applied the nRF distance between the obtained and the reference trees as a measure of accuracy.

We simulated five sets of 33 genomes, each with different extents of HGT as determined by the mean number of HGT events per iteration (*l* = 0, 250, 500, 750, and 1,000; *l* is the number of HGT events attempted in the set at each iteration of the simulation process of genome evolution; for details, see **Methods**). The tools, AFKS (Markov measure, with a word length of *k* = 12) and mash (*k* = 17-24) achieved the highest general accuracy (**Figure 2**) by obtaining the lowest average nRF (0.05 ± 0.05) and perfect topological agreement with the reference trees at the two lowest frequencies of simulated HGT (*l* = 0 and 250; Additional file 1: **Table S11**). As expected, for most AF methods, the accuracy of phylogenetic inference declines with an increase in the extent of HGT. Nevertheless, the seven best-performing software applications — AFKS, mash, CAFE, alfpy, FFP, jD2Stat, and ALFRED-G [57] — were capable of reconstructing the reference tree with little incongruence at almost all HGT frequency levels (nRF ≤ 0.1 at *l* ≤ 750), except for the highest frequencies of HGT simulated, where the nRF distance was in the range of 0.13-0.17 (Additional file 1: **Table S11**). Interestingly, the basic AF distance measures (Euclidean, Manhattan, Canberra and LCC distances) implemented in alfpy achieve a lower average nRF (0.07 ± 0.06) and minimum nRF at a higher HGT frequency level (nRF = 0.13) than AF tools designed for phylogenetic reconstruction of whole genomes (co-phylog, FSWM, Multi-SpaM and kr), which surprisingly were relatively inaccurate (nRF > 0.2 for different values of *l*). As has been reported before [53], the accuracy of kr generally increased (nRF: from 0.73 to 0.33) with increasing *l*.

To assess the performance of AF methods with real-world sequence data, we first used a reference supertree of 27 genomes of *E. coli* and *Shigella* that was generated based on thousands of single-copy protein trees [53–55]. For this data set, the tools designed for whole-genome phylogenetics achieved lower nRF values than did basic AF distance measures; eleven tools for whole-genome phylogenetics occupied the first six positions in the ranking list (**Figure 2**). Three such methods — andi, co-phylog and phylonium — achieved the highest accuracy (**Figure 2**), with a minimum nRF of 0.08 (Additional file 1: **Table S12**). The andi and co-phylog tools yielded topologically equivalent trees that were very similar to the reference tree, misplacing only two closely related *E. coli* strains in the D and B1 reference groups (Additional file 2: **Figure S7**), while phylonium showed two minor topological differences in *E. coli* reference group D (Additional file 2: **Figure S8**). Most AF measures implemented in AFKS, alfpy and CAFE were ranked at the 10^th^ position (**Figure 2**) and led to reconstruction of inaccurate species trees where half of the bipartitions were not present in the reference tree (nRF = 0.5). Interestingly, the opposite result was obtained for phylogenetic inference of 8 *Yersinia* genomes, where almost all basic measures (42 tool variants) recovered the reference tree topology (nRF = 0) while whole-genome phylogenetic tools obtained relatively incongruent trees (nRF > 0.2) compared to the reference (**Figure 2**, Additional file 1: **Table S13**).

## DISCUSSION

We have addressed key challenges in assessing methods for AF sequence comparison by automating the application of multiple AF methods to a range of reference data sets. This automated approach critically benefits from extensive work described in the previous section to identify optimal parameter values for all combinations of methods and data sets. Finally, the resulting open platform for a standardized evaluation of new methods is provided with an interactive web-based interface and a reporting functionality designed to ensure reproducibility. We believe that the uniform framework for testing AF algorithms with common data sets and procedures will be beneficial to both developers and users of these methods. The benchmarking results will guide users in choosing the most effective tool tailored to their project needs and for finding optimal parameter settings, improving the quality of their studies and results. For developers, the interactive platform speeds up benchmarking and provides reference data sets, on which new AF methods can be compared to existing approaches.

Our results showed that no single method performed best across all the data sets tested. Nevertheless, some tools were among the top five performers more often than others. For example, when considering genomic-scale benchmarks, encompassing 8 datasets from the whole-genome phylogeny and horizontal gene transfer categories, the tools developed for genomic comparisons were among the top-5-performing tools: mash (8 times), co-phylog and Skmer (7 times), FFP (6 times), and FSWM/Read-SpaM (5 times; **Figure 2**). Since mash is the only method that is placed among the top five-best performing tools on all genome-scale benchmarking data sets, it is particularly well suited for genome sequence comparisons, regardless of the phylogenetic range and technology that were used to obtain the data (e.g., short reads or assembled contigs). Most AF approaches (14 out of 21 software applications or, more specifically, 56 out of 68 tool variants) performed particularly well — although not perfectly — in phylogenetic inference of mitochondrial genomes from different fish species, yielding trees generally consistent (nRF < 0.1) with the reference phylogeny (**Figure 2**, Additional file 1: **Table S6**). However, our results on whole-genome sequence comparison for prokaryotes and eukaryotes show significant decrease in performance of tested AF tools. Thus, novel AF methods should not be benchmarked with mitochondrial sequences alone. Considering the evolutionary and structural relationships among the protein sequences and inferred gene trees, we were surprised by the highest performance of very simple AF distance measures implemented in AFKS and alfpy (i.e., intersection, simratio, Kulczynski, Bray-Curtis, Google, Canberra, Squared_chord, chi_squared, and Manhattan). Overall, methods based on conventional statistics performed better than approaches using more complex statistics such as state-of-the-art D_2_-related metrics implemented in jD2Stat (D_2_^S^, D_2_*, and D_2_^n^) and AFKS (D_2_^z^, D_2_*, and D_2_^S^), the Markov metric in AFSK (sim_mm, rr_k_r, and markov), and the N_2_ metric in AFKS (n_2_r) (Additional file 1: **Table S14**). Interestingly, the basic Canberra distance implemented in alfpy is the most effective distance measure in recognizing functionally related regulatory sequences (Additional file 1: **Table S5**), greatly exceeding the D_2_S and D_2_* statistics from CAFE and jD2Stat.

Another surprising observation in our study is that different implementations of the same AF algorithm, run with the same input parameter values, can deliver different results. For example, two implementations of the Canberra distance from AFKS and alfpy achieve different performances in almost all data sets (**Figure 2**). The discrepancy in the Canberra distance with a word length of *k* = 2 between the two tools is apparent for the CRM data set, where AFKS--*Canberra* obtained a performance score of 54, while alfpy--*Canberra* had a performance score of 74, which was the highest performance score among the tools that we evaluated (Additional file 1: **Table S5**; see the **Methods** section for the definition of “performance score”). The differences observed were due to the different methods of sequence data preprocessing applied by the two tools — alfpy projects sequences into a vector of *k*-mer frequencies, whereas AFKS represents sequences as *k*-mer count vectors with the inclusion of pseudocounts. This sequence data preprocessing in alfpy and AFKS has the highest impact on the performance of methods based on the Canberra distance in the case of nucleotide data sets of regulatory elements, whole genomes of plants and simulated genomes that underwent HGT (Additional file 2: **Figure S9**). For other data sets, the same distance measures in alfpy and AFKS, run on common word lengths, produce results with very similar performances, and the observed differences between the tools in this study are the results of different ranges of *k*. Similarly, the *D_2_\** and *D_2_^S^* metrics implemented in AFKS, CAFE and jD2Stat produce slightly different results.

When assessing the accuracy of AF methods in inferring phylogenetic relationships, we compared the inferred phylogenetic tree topologies to trusted reference tree topologies. However, the assumption that evolutionary relationships are generally tree-like is known to be unrealistic because genome evolution is shaped by both vertical and lateral processes [55,58,59]. Although the signal of vertical descent (e.g., for ribosomal rRNAs) can be described adequately using a phylogenetic tree, horizontal transfer of genetic material between different taxa and genome rearrangements can obscure this signal. A classic example involves the *Yersinia* genomes, which are well-known to have undergone extensive structural rearrangements [56]. We have shown in this study that reconstructing phylogenetic trees of these taxa from whole-genome sequences is difficult with AF methods. The same is true for more conventional approaches that are based on MSA [56], and finding a trusted reference tree for these taxa has been problematic. In such cases, a non-tree-like network representation of genome evolution is more appropriate. Recent studies [60,61] have demonstrated the scalability and applicability of AF methods to quickly infer networks of relatedness among microbial genomes. Although we did not consider networks in this study, the curated benchmarking data sets can be easily extended to AF phylogenetic analysis beyond a tree-like structure in the future.

We acknowledge that the presented data sets do not cover all possible applications of AF tools. The data sets include only the most typical sequence comparison tasks, where all-versus-all sequence comparisons need to be computed. Although the AF project is extendable and new data sets can be seamlessly added in the future, for more specific applications such as orthology prediction, genome assembly, RNA-seq aligners or metagenomics analyses, we recommend using other web-based benchmarking services developed for these purposes [37,62–65]. Nevertheless, AFproject can be used to evaluate any sequence comparison tool — not necessarily AF — that produces dissimilarity scores between sequence pairs. Since similarity scores can be easily converted into dissimilarity scores, our benchmarking system can also be used to evaluate methods that generate similarity scores, e.g., alignment scores. We thus invite developers and users of sequence comparison methods to submit and evaluate their results with the AFproject benchmarking platform. The ability to rapidly, objectively and collaboratively compare computational methods for sequence comparison should be beneficial for all fields of DNA and RNA sequence analysis, regardless of whether the analysis is alignment-based or alignment-free.

## METHODS

### Data sets

Twelve sequence data sets were used to evaluate AF methods across five research areas (**Table 1**).

#### Protein homology

The reference data sets of protein family members sharing a high (≥40%) and low (<40%) sequence identity were constructed based on two sections of the SCOPe database v. 2.07 [39], namely, ASTRAL95 and ASTRAL40 v. 2.07 [66], respectively. The SCOPe database provides a structural classification of proteins at four levels: classes, folds, superfamilies and families. According to previous studies [3,6], the ASTRAL data sets were subsequently trimmed to exclude sequences with unknown amino acids and families with fewer than 5 proteins and included only the four major classes (i.e., α, β, α/β, and α+β). To minimize the requirements for AF method submission related to performing all-versus-all sequence comparisons and uploading the output to the AFproject server, we further reduced the data sets by randomly selecting only two protein members in each family. As ASTRAL95 also contains protein family members sharing a sequence identity lower than 40%, the Needleman-Wunsch alignment was performed (using needle software in the EMBOSS package [67]) to select proteins with a sequence identity ≥ 40% to acquire a reference data set of proteins with high sequence identity.

#### Gene trees

Reference trees and corresponding protein sequences of eleven gene families were downloaded from SwissTree release 2017.0 (https://swisstree.vital-it.ch/): Popeye domain-containing protein family (49 genes), NOX ‘ancestral-type’ subfamily NADPH oxidases (54 genes), V-type ATPase beta subunit (49 genes), Serine incorporator family (115 genes), SUMF family (29 genes), Ribosomal protein S10/S20 (60 genes), Bambi family (42 genes), Asterix family (39 genes), Cited family (34 genes), Glycosyl hydrolase 14 family (159 genes), and Ant transformer protein (21 genes).

#### Gene regulatory elements

The data set of CRMs known to regulate expression in the same tissue and/or developmental stage in fly or human was obtained from Kantorovitz et al. [4]. The data set was specifically selected to test the capacity of AF measures to identify functional relationships among regulatory sequences (e.g., enhancers or promoters). The data set contains 185 CRM sequences taken from *D. melanogaster* — blastoderm-stage embryo (*n* = 82), eye (*n* = 17), peripheral nervous system (*n* = 23), and tracheal system (*n* = 9) — and *Homo sapiens* — HBB complex (*n* = 17), liver (*n* = 9) and muscle (*n* = 28).

#### Genome-based phylogeny

The sequences of 25 whole mitochondrial genomes of fish species from the suborder Labroidei and the species tree were taken from Fischer et al. [48]. The set of 29 *E. coli* genome sequences was originally compiled by Yin and Jin [21] and has been used in the past by other groups to evaluate AF programs [22,23,68]. Finally, the set of 14 plant genomes is from Hatje et al. [69]. This set was also used in the past to evaluate AF methods. To simulate unassembled reads from these data sets, we used the program ART [70].

#### Horizontal gene transfer

The 27 *E. coli* and *Shigella* genomes, and the 8 *Yersinia* genomes, were taken from Bernard et al. [53]. We used EvolSimulator [71] to simulate HGT in microbial genomes, adopting an approach similar to that described in Bernard et al. [53]. Each set of genomes was simulated under a birth-and-death model at speciation rate = extinction rate = 0.5. The number of genomes in each set was allowed to vary from 25 to 35, with each containing 2,000–3,000 genes 240–1,500 nucleotides long. HGT receptivity was at set at a minimum of 0.2, mean of 0.5 and maximum of 0.8, with a mutation rate *m* = 0.4–0.6 and a number of generations *i* = 5,000. The varying extent of HGT was simulated using the mean number of HGT events attempted per iteration *l* = 0, 250, 500, 750 and 1000, and divergence factor *d* = 2,000 (transferred genes that are of high sequence divergence, i.e., >2,000 iterations apart, will not be successful). All other parameters in this simulation followed Beiko et al. [71].

### Alignment-free tools

AAF [72] reconstructs a phylogeny directly from unassembled next-generation sequencing reads. Specifically, AAF calculates the Jaccard distance between sets of *k*-mers of two samples of short sequence reads. This distance between samples or species is based on the estimate of the rate parameter from a Poisson process for a mutation occurring at a single nucleotide. The phylogeny is constructed using weighted least squares with weights proportional to the expected variance of the estimated distances. AAF provides features for correcting tip branches and bootstrapping of the obtained phylogenetic trees, directly addressing the problems of sequencing error and incomplete coverage. AFKS [32] is a package for calculating 33 *k*-mer-based dissimilarity/distance measures between nucleotide or protein sequences. AFKS categorizes the measures into nine families: Minkowski (e.g., Euclidean), Mismatch (e.g., Jaccard), Intersection (e.g., Kulczynski), D2 (e.g., D2s), Squared Chord (e.g., Hellinger), Inner Product (e.g., normalized vectors), Markov (e.g., SimMM), Divergence (e.g., KL Conditional), and Others (e.g., length difference). The tool determines the optimal *k*-mer size for given input sequences and calculates dissimilarity/distance measures between *k*-mer counts that include pseudocounts (adding 1 to each *k*-mer count). The obtained distance is standardized to between 0 and 1.

alfpy [3] provides 38 AF dissimilarity measures with which to calculate distances among given nucleotide or protein sequences. The tool includes 25 *k*-mer-based measures (e.g., Euclidean, Minkowski, Jaccard, and Hamming), eight information-theoretic measures (e.g., Lempel–Ziv complexity and normalized compression distance), three graph-based measures, and two hybrid measures (e.g., Kullback-Leibler divergence and W-metric). alfpy is also available as a web application and Python package. In this study, the results based on 14 dissimilarity measures are evaluated.

ALFRED-G [57] uses an efficient algorithm to calculate the length of maximal *k*-mismatch common substrings between two sequences. Specifically, to measure the degree of dissimilarity between two nucleic acid or protein sequences, the program calculates the length of maximal word pairs — one word from each of the sequences — with up to *k* mismatches.

andi [22] estimates phylogenetic distances between genomes of closely related species by identifying pairs of maximal unique word matches a certain distance from each other and on the same diagonal in the comparison matrix of two sequences. Such word matches can be efficiently found using enhanced suffix arrays. The tool then uses these gap-free alignments to estimate the number of substitutions per position.

CAFE [34] is a package for efficient calculation of 28 AF dissimilarity measures, including 10 conventional measures based on *k*-mer counts, such as Chebyshev, Euclidean, Manhattan, uncentered correlation distance, and Jensen-Shannon divergence. It also offers 15 measures based on the presence/absence of *k*-mers, such as Jaccard and Hamming distances. Most importantly, it provides a fast calculation of background-adjusted dissimilarity measures including CVTree, d2star and d2shepp. CAFE allows for both assembled genome sequences and unassembled next-generation sequencing shotgun reads as inputs. However, it does not deal with amino acid sequences. In this study, the results based on CVTree, d2star and d2shepp are evaluated.

co-phylog [21] estimates evolutionary distances among assembled or unassembled genomic sequences of closely related microbial organisms. The tool finds short, gap-free alignments of a fixed length and consisting of matching nucleotide pairs only, except for the middle position in each alignment, where mismatches are allowed. Phylogenetic distances are estimated from the fraction of such alignments for which the middle position is a mismatch.

EP-sim [73] computes an AF distance between nucleotide or amino acid sequences based on entropic profiles [74,75]. The entropic profile is a function of the genomic location that captures the importance of that region with respect to the whole genome. For each position, it computes a score based on the Shannon entropies of the word distribution and variable-length word counts. EP-sim estimates a phylogenetic distance, similar to *D_2_*, by summing the entropic profile scores over all positions, or similar to *D_2_\**, with the sum of normalized entropic profile scores.

FFP [33,38] estimates the distances among nucleotide or amino acid sequences. The tool calculates the count of each *k*-mer and then divides the count by the total count of all *k*-mers to normalize the counts into frequencies of a given sequence. This process leads to the conversion of each sequence into its feature frequency profile (FFP). The pairwise distance between two sequences is then calculated by the Jensen-Shannon divergence between their respective FFPs.

FSWM [24] estimates the phylogenetic distance between two DNA sequences. The program first defines a fixed binary pattern *P* of length *l* representing “match positions” and “don’t care positions”. Then, it identifies all “<u>Spa</u>ced-word <u>M</u>atches” (*SpaM*) w.r.t. *P*, i.e., gap-free local alignments of the input sequences of length *l*, with matching nucleotides at the “match positions” of *P* and possible mismatches at the “don’t care” positions. To estimate the distance between two DNA sequences, *SpaM*s with low overall similarity are discarded, and the remaining *SpaM*s are used to estimate the distance between the sequences, based on the mismatch ratio at the “don’t care” positions. There is a version of FSWM that can compare sets of unassembled sequencing reads to each other called *Read-SpaM* [76].

jD2Stat [35] utilizes a series of *D_2_* statistics [15,16] to extract *k*-mers from a set of biological sequences and generate pairwise distances for each possible pair as a matrix. For each sequence set, we generated distance matrices (at the defined *k*; Additional file 1: **Table S1**), each using *D_2_^S^* (D2S; exact *k*-mer counts normalized based on the probability of occurrences of specific *k*-mers), *D_2_\** (d2St; similar to *D_2_^S^* but normalized based on means and variance), and *D_2_^n^* (d2n; extension of *D_2_* that expands each word *w* recovered in the sequences to its neighborhood *n*, i.e., all possible *k*-mers with *n* number of wildcard residues, relative to *w*).

kmacs [18] compares two DNA or protein sequences by searching for the longest common substrings with up to *k* mismatches. More precisely, for each position *i* in one sequence, the program identifies the longest pair of substrings with up to *k* mismatches, starting at *i* in the first sequence and somewhere in the second sequence. The average length of these substring pairs is then used to define the distance between the sequences.

kr [49] estimates the evolutionary distance between genomes by calculating the number of substitutions per site. The estimator for the rate of substitutions between two unaligned sequences depends on a mathematical model of DNA sequence evolution and average shortest unique substring (shustring) length.

kSNP3 [77] identifies single nucleotide polymorphisms (SNPs) in a set of genome sequences without the need for genome alignment or a reference genome. The tool defines a SNP locus as the *k*-mers surrounding a central SNP allele. kSNP3 can analyze complete genomes, draft genomes at the assembly stage, genomes at the raw reads stage, or any combination of these stages. Based on the identified SNPs, kSNP3.0 estimates phylogenetic trees by parsimony, neighbor-joining and maximum-likelihood methods and reports a consensus tree with the number of SNPs unique to each node.

kWIP [78] estimates genetic dissimilarity between samples directly from next-generation sequencing data without the need for a reference genome. The tool uses the weighted inner product (WIP) metric, which aims at reducing the effect of technical and biological noise and elevate the relevant genetic signal by weighting *k*-mer counts by their informational entropy across the analysis set. This procedure downweights *k*-mers that are typically uninformative (highly abundant or present in very few samples).

LZW-Kernel [79] classifies protein sequences and identifies remote protein homology via a convolutional kernel function. LZW-Kernel exploits code blocks detected by the universal Lempel-Ziv-Welch (LZW) text compressors and then builds a kernel function out of them. LZW-Kernel provides a similarity score between sequences from 0 to 1, which can be directly used with support vector machines (SVMs) in classification problems. LZW-Kernel can also estimate the distance between protein sequences using normalized compression distances (LZW-NCD).

mash [9] estimates the evolutionary distance between nucleotide or amino acid sequences. The tool uses the MinHash algorithm to reduce the input sequences to small ‘sketches’, which allow fast distance estimations with low storage and memory requirements. To create a ‘sketch’, each *k*-mer in a sequence is hashed, which creates a pseudorandom identifier (hash). By sorting these hashes, a small subset from the top of the sorted list can represent the entire sequence (min-hashes). Two sketches are compared to provide an estimate of the Jaccard index (i.e., the fraction of shared hashes) and the Mash distance, which estimates the rate of sequence mutation under an evolutionary model.

Multi-SpaM [23], similarly to FSWM, starts with a binary pattern *P* of length *l* representing “match positions” and “don’t care positions”. It then searches for four-way <u>Spa</u>ced-word <u>M</u>atches (*SpaMs*) w.r.t. *P*, i.e., local gap-free alignments of length *l* involving four sequences each and with identical nucleotides at the “match positions” and possible mismatches at the “don’t care positions”. Up to 1,000,000 such multiple SpaMs with a score above some threshold are randomly sampled, and a quartet tree is calculated for each of them with RAxML [80]. The program *Quartet Max-Cut* [81] is used to calculate a final tree of all input sequences from the obtained quartet trees.

phylonium [51] estimates phylogenetic distances among closely related genomes. The tool selects one reference from a given set of sequences and finds matching sequence segments of all other sequences against this reference. These long and unique matching segments (anchors) are calculated using an enhanced suffix array. Two equidistant anchors constitutes homologous region, in which SNPs are counted. With the analysis of SNPs, phylonium estimates the evolutionary distances between the sequences.

RTD-Phylogeny [82] computes phylogenetic distances among nucleotide or protein sequences based on the time required for the reappearance of *k*-mers. The time refers to the number of residues in successive appearance of particular *k*-mers. Thus, the occurrence of each *k*-mer in a sequence is calculated in the form of a return time distribution (RTD), which is then summarized using the mean (µ) and standard deviation (σ). As a result, each sequence is represented in the form of a numeric vector of size 2·4^k^ containing the µ and σ of 4^k^ RTDs. The pairwise distance between sequences is calculated using Euclidean distance.

Skmer [52] estimates phylogenetic distances between samples of raw sequencing reads. Skmer runs mash [9] internally to compute the *k*-mer profile of genome skims and their intersection, and estimates the genomic distances by correcting for the effect of low coverage and sequencing error. The tool can estimate distances between samples with high accuracy from low-coverage and mixed-coverage genome skims with no prior knowledge of the coverage or the sequencing error.

Slope-SpaM [83] estimates the phylogenetic distance between two DNA sequences by calculating the number *N_k_* of *k*-mer matches for a range of values of *k*. The distance between the sequences can then be accurately estimated from the *slope* of a certain function that depends on *N_k_*. Instead of exact word matches, the program can also use *SpaMs* w.r.t. a predefined binary pattern of “match positions” and “don’t care positions”.

spaced [84–86] is similar to previous methods that compare the *k*-mer composition of DNA or protein sequences. However, the program uses so-called “spaced words” instead of *k*-mers. For a given binary pattern *P* of length *l* representing “match positions” and “don’t care positions”, a spaced word w.r.t. *P* is a word of length *l* with nucleotide or amino acid symbols at the “match positions” and “wildcard characters” at the “don’t care positions”. The advantage of using spaced words instead of exact *k*-mers is that the obtained results are statistically more stable. This idea has been previously proposed for database searching [87,88]. The original version of Spaced [84] used the Euclidean or Jensen-Shannon [89] distance to compare the spaced-word composition of genomic sequences. By default, the program now uses a distance measure introduced by Morgenstern et al. 2015 [86] that estimates the number of substitutions per sequence position.

Underlying Approach [90] estimates phylogenetic distances between whole genomes using matching statistics of common words between two sequences. The matching statistics are derived from a small set of independent subwords with variable lengths (termed *irredundant common subwords*). The dissimilarity between sequences is calculated based on the length of the longest common subwords, such that each region of genomes contributes only once, thus avoiding counting shared subwords multiple times (i.e., subwords occurring in genomic regions covered by other more significant subwords are discarded).

### Benchmarks

#### Evaluation of structural and evolutionary relationships among proteins

To test the capacity of AF distance measures to recognize SCOPe relationships (i.e., family, superfamily, fold, and class), we used a benchmarking protocol from previous studies [6], [3]. Accordingly, the benchmarking procedure takes the distances between all sequence pairs present in the data set file. The distances between all protein pairs are subsequently sorted from minimum to maximum (i.e., from the maximum to minimum similarity). The comparative test procedure is based on a binary classification of each protein pair, where 1 corresponds to the two proteins sharing the same group in the SCOPe database and 0 corresponds to other outcomes. The group can be defined at one of the four different levels of the database (family, superfamily, fold, and class), exploring the hierarchical organization of the proteins in that structure. Therefore, each protein pair is associated with four binary classifications, one for each level. At each SCOPe level, ROC curves and AUC values computed in scikit-learn [91] are obtained to give a unique number of the relative accuracy of each metric and level according to the SCOP classification scheme. The overall assessment of method accuracy is an average of AUC values across all four SCOPe levels.

#### Evaluation of functionally related regulatory sequences

To test how well AF methods can capture the similarity between sequences with similar functional roles, we used the original benchmarking protocol introduced by Kantorovitz et al. [4]. Briefly, a set of CRMs known to regulate expression in the same tissue and/or developmental stage is taken as the ‘positive’ set. An equally sized set of randomly chosen noncoding sequences with lengths matching the CRMs is taken as the ‘negative’ set. Each pair of sequences in the positive set is compared, as is each pair in the negative set. The test evaluates if functionally-related CRM sequence pairs (from the positive half) are better scored by a given AF tool (i.e., have lower distance/dissimilarity values) than unrelated pairs of sequences (from the negative half). This procedure is done by sorting all pairs, whether they are from the positive set or the negative set, in one combined list and then counting how many of the pairs in the top half of this list are from the positive set. The overall assessment of method accuracy is the weighted average of the positive pairs across all seven subsets.

#### Evaluation of phylogenetic inference

The accuracy of AF methods for data sets from three categories — protein homology, genome-based phylogeny and horizontal gene transfer — was evaluated by a comparison of topology between the method’s tree and the reference tree. The pairwise sequence distances obtained by the AF method were used as input for the neighbor-joining algorithm (fneighbor in the EMBOSS package [67], version: EMBOSS:6.6.0.0 PHYLIPNEW:3.69.650) to generate the corresponding method tree. To assess the degree of topological (dis)agreement between the inferred and reference trees, we calculated the nRF distance [36] using the Tree.compare function in the ETE3 [92] toolkit for phylogenetic trees with the option unrooted=True. When nRF = 0, the test and reference topologies are identical, implying the highest accuracy for the method. Conversely, at nRF = 1, no bipartition in the reference is recovered.

#### Performance summary criteria

Figure 2 shows the color-coded performance of the evaluated AF methods across 12 reference data sets.

#### Performance score

For our benchmarking data sets, we use different measures to assess the performance of each method for a given data set, for example, nRF or AUC. To make our benchmarking results from different data sets comparable, we converted these measures to a performance score with values between 0 and 100. For the protein sequence classification data sets, this score is defined as AUC × 100; for data sets from gene trees, genome-based phylogeny and horizontal gene transfer categories, we define the performance score as (1 – nRF) × 100. For the regulatory elements data set, the performance score is already a number between 0 and 100, namely, the weighted average performance across seven data subsets. Moreover, we define an *overall performance score* (Additional file 1: **Table S14**) that assesses each method across the data sets and that also takes values between 0 and 100. For a given method, we calculate revised scores for each data set, on which the method was tested as (*S* – *min_score*) / (*max_score* – *min_score*) × 100, where *S* is the performance score obtained by the method and *min_score* and *max_score* are the minimum and maximum scores obtained with all methods for a given data set, respectively. This way, the best-performing method in a given data set receives a score of 1, and the worst performer receives a score of 0. The overall performance is an average of the revised scores across the data sets on which the given method was tested.

## ADDITIONAL FILES

**Additional file 1:** Supplementary Tables S1-S14

**Additional file 2:** Supplementary Figures S1-S9

## ACKNOWLEDGMENTS

We thank Svenja Dörrer for providing benchmarking data sets of unassembled sequencing reads. We also thank the tools developers B. Haubold, F. Klötzl, P. Kolekar, S. Mirarab, S. Sarmashghi, A. Phillippy, and H. Yi for recommending analysis parameters and helpful discussions. Computations were performed at Poznan Supercomputing and Networking Center (PSNC).

## AVAILABILITY OF DATA AND MATERIALS

All data sets and results discussed in the paper are freely available from our website (http://afproject.org). The source code of the AFproject service is available under an open source license (Mozilla Public License Version 2.0) at https://github.com/afproject-org/afproject.

## AUTHOR CONTRIBUTIONS

A.Z. and W.M.K. conceived the project. B.M., A.K.L., CX.C, G.B., A.Z. and W.M.K contributed the reference data sets. A.Z. and W.M.K. designed and implemented the benchmarking service. A.Z., H.Z.G., B.T.J., G.B., C.L., K.T., T.D., J.C., M.C., S.K, S.R. and M.S.W. contributed the benchmarking results. A.Z., B.M., F.S., S.V. and W.M.K. analyzed the results. A.Z., B.M., S.V., J.S.A., CX.C., H.Z.G., J.C. and W.M.K. prepared the manuscript, with feedback from all other coauthors. W.M.K. coordinated the project.

## Ethics approval and consent to participate

Not applicable.

## REFERENCES

1. Altschul SF, Gish W, Miller W, Myers EW, Lipman DJ. Basic local alignment search tool. J Mol Biol. 1990;215:403–10.

2. Thompson JD, Higgins DG, Gibson TJ. CLUSTAL W: improving the sensitivity of progressive multiple sequence alignment through sequence weighting, position-specific gap penalties and weight matrix choice. Nucleic Acids Res. 1994;22:4673–80.

3. Zielezinski A, Vinga S, Almeida J, Karlowski WM. Alignment-free sequence comparison: benefits, applications, and tools. Genome Biol. 2017;18:186.

4. Kantorovitz MR, Robinson GE, Sinha S. A statistical method for alignment-free comparison of regulatory sequences. Bioinformatics. 2007;23:i249–55.

5. Ivan A, Halfon MS, Sinha S. Computational discovery of cis-regulatory modules in Drosophila without prior knowledge of motifs. Genome Biol. 2008;9:R22.

6. Vinga S, Gouveia-Oliveira R, Almeida JS. Comparative evaluation of word composition distances for the recognition of SCOP relationships. Bioinformatics. 2004;20:206–15.

7. Terrapon N, Weiner J, Grath S, Moore AD, Bornberg-Bauer E. Rapid similarity search of proteins using alignments of domain arrangements. Bioinformatics. 2014;30:274–81.

8. Cong Y, Chan Y-B, Ragan MA. A novel alignment-free method for detection of lateral genetic transfer based on TF-IDF. Sci Rep. 2016;6:30308.

9. Ondov BD, Treangen TJ, Melsted P, Mallonee AB, Bergman NH, Koren S, et al. Mash: fast genome and metagenome distance estimation using MinHash. Genome Biol. 2016;17:132.

10. Fox GE, Magrum LJ, Balch WE, Wolfe RS, Woese CR. Classification of methanogenic bacteria by 16S ribosomal RNA characterization. Proc Natl Acad Sci U S A. 1977;74:4537–41.

11. Vinga S, Almeida J. Alignment-free sequence comparison--a review. Bioinformatics. 2003;19:513–23.

12. Jun S-R, Sims GE, Wu GA, Kim S-H. Whole-proteome phylogeny of prokaryotes by feature frequency profiles: An alignment-free method with optimal feature resolution. Proc Natl Acad Sci U S A. 2010;107:133–8.

13. Sims GE, Kim S-H. Whole-genome phylogeny of Escherichia coli/Shigella group by feature frequency profiles (FFPs). Proc Natl Acad Sci U S A. 2011;108:8329–34.

14. Blaisdell BE. A measure of the similarity of sets of sequences not requiring sequence alignment. Proc Natl Acad Sci U S A. 1986;83:5155–9.

15. Reinert G, Chew D, Sun F, Waterman MS. Alignment-free sequence comparison (I): statistics and power. J Comput Biol. 2009;16:1615–34.

16. Wan L, Reinert G, Sun F, Waterman MS. Alignment-free sequence comparison (II): theoretical power of comparison statistics. J Comput Biol. 2010;17:1467–90.

17. Ulitsky I, Burstein D, Tuller T, Chor B. The average common substring approach to phylogenomic reconstruction. J Comput Biol. 2006;13:336–50.

18. Leimeister C-A, Morgenstern B. Kmacs: the k-mismatch average common substring approach to alignment-free sequence comparison. Bioinformatics. 2014;30:2000–8.

19. Yang L, Zhang X, Fu H, Yang C. An estimator for local analysis of genome based on the minimal absent word. J Theor Biol. 2016;395:23–30.

20. Yang L, Zhang X, Zhu H. Alignment free comparison: similarity distribution between the DNA primary sequences based on the shortest absent word. J Theor Biol. 2012;295:125–31.

21. Yi H, Jin L. Co-phylog: an assembly-free phylogenomic approach for closely related organisms. Nucleic Acids Res. 2013;41:e75.

22. Haubold B, Klötzl F, Pfaffelhuber P. andi: fast and accurate estimation of evolutionary distances between closely related genomes. Bioinformatics. 2015;31:1169–75.

23. Dencker T, Leimeister C-A, Gerth M, Bleidorn C, Snir S, Morgenstern B. Multi-SpaM: A Maximum-Likelihood Approach to Phylogeny Reconstruction Using Multiple Spaced-Word Matches and Quartet Trees. Lecture Notes in Computer Science. 2018. p. 227–41.

24. Leimeister C-A, Sohrabi-Jahromi S, Morgenstern B. Fast and accurate phylogeny reconstruction using filtered spaced-word matches. Bioinformatics. 2017;33:971–9.

25. Leimeister C-A, Schellhorn J, Dörrer S, Gerth M, Bleidorn C, Morgenstern B. Prot-SpaM: fast alignment-free phylogeny reconstruction based on whole-proteome sequences. Gigascience [Internet]. 2019;8. Available from: https://doi.org/10.1093/gigascience/giy148

26. Almeida JS, Carrico JA, Maretzek A, Noble PA, Fletcher M. Analysis of genomic sequences by Chaos Game Representation. Bioinformatics. 2001;17:429–37.

27. Jeffrey HJ. Chaos game representation of gene structure. Nucleic Acids Res. 1990;18:2163–70.

28. Yau SS-T, -T. Yau SS, Yu C, He R. A Protein Map and Its Application. DNA Cell Biol. 2008;27:241–50.

29. Yin C, Yau SS-T. An improved model for whole genome phylogenetic analysis by Fourier transform. J Theor Biol. 2015;382:99–110.

30. Vinga S. Information theory applications for biological sequence analysis. Brief Bioinform. 2014;15:376–89.

31. Almeida JS. Sequence analysis by iterated maps, a review. Brief Bioinform. 2014;15:369–75.

32. Luczak BB, James BT, Girgis HZ. A survey and evaluations of histogram-based statistics in alignment-free sequence comparison. Brief Bioinform [Internet]. 2017; Available from: http://dx.doi.org/10.1093/bib/bbx161

33. Sims GE, Jun S-R, Wu GA, Kim S-H. Alignment-free genome comparison with feature frequency profiles (FFP) and optimal resolutions. Proc Natl Acad Sci U S A. 2009;106:2677–82.

34. Lu YY, Tang K, Ren J, Fuhrman JA, Waterman MS, Sun F. CAFE: aCcelerated Alignment-FrEe sequence analysis. Nucleic Acids Res. 2017;45:W554–9.

35. Chan CX, Bernard G, Poirion O, Hogan JM, Ragan MA. Inferring phylogenies of evolving sequences without multiple sequence alignment. Sci Rep. 2014;4:6504.

36. Robinson DF, Foulds LR. Comparison of phylogenetic trees. Math Biosci. 1981;53:131–47.

37. Altenhoff AM, Boeckmann B, Capella-Gutierrez S, Dalquen DA, DeLuca T, Forslund K, et al. Standardized benchmarking in the quest for orthologs. Nat Methods. 2016;13:425–30.

38. Choi J, Kim S-H. A genome Tree of Life for the Fungi kingdom. Proc Natl Acad Sci U S A. 2017;114:9391–6.

39. Fox NK, Brenner SE, Chandonia J-M. SCOPe: Structural Classification of Proteins--extended, integrating SCOP and ASTRAL data and classification of new structures. Nucleic Acids Res. 2014;42:D304–9.

40. Wu TJ, Burke JP, Davison DB. A measure of DNA sequence dissimilarity based on Mahalanobis distance between frequencies of words. Biometrics. 1997;53:1431–9.

41. Hide W, Burke J, Davison DB. Biological evaluation of d2, an algorithm for high-performance sequence comparison. J Comput Biol. 1994;1:199–215.

42. Rokas A, Williams BL, King N, Carroll SB. Genome-scale approaches to resolving incongruence in molecular phylogenies. Nature. 2003;425:798–804.

43. Boeckmann B, Dylus D, Moretti S, Altenhoff A, Train C-M, Kriventseva E, et al. Taxon sampling unequally affects individual nodes in a phylogenetic tree: consequences for model gene tree construction in SwissTree [Internet]. 2017. Available from: http://dx.doi.org/10.1101/181966

44. Dai Q, Yang Y, Wang T. Markov model plus k-word distributions: a synergy that produces novel statistical measures for sequence comparison. Bioinformatics. 2008;24:2296–302.

45. Chan CX, Ragan MA. Next-generation phylogenomics. Biol Direct. BioMed Central; 2013;8:3.

46. Haubold B. Alignment-free phylogenetics and population genetics. Brief Bioinform. 2014;15:407–18.

47. Li M, Badger JH, Chen X, Kwong S, Kearney P, Zhang H. An information-based sequence distance and its application to whole mitochondrial genome phylogeny. Bioinformatics. 2001;17:149–54.

48. Fischer C, Koblmüller S, Gülly C, Schlötterer C, Sturmbauer C, Thallinger GG. Complete mitochondrial DNA sequences of the threadfin cichlid (Petrochromis trewavasae) and the blunthead cichlid (Tropheus moorii) and patterns of mitochondrial genome evolution in cichlid fishes. PLoS One. 2013;8:e67048.

49. Haubold B, Pfaffelhuber P, Domazet-Loso M, Wiehe T. Estimating Mutation Distances from Unaligned Genomes. J Comput Biol. 2009;16:1487–500.

50. Lin J, Adjeroh DA, Jiang B-H, Jiang Y. K2 and K2*: efficient alignment-free sequence similarity measurement based on Kendall statistics. Bioinformatics. 2018;34:1682–9.

51. Fabian K, Haubold B. Phylonium – fast and accurate estimation of evolutionary distances [Internet]. GitHub. [cited 2019 Feb 10]. Available from: https://github.com/kloetzl/phylonium

52. Sarmashghi S, Bohmann K, P Gilbert MT, Bafna V, Mirarab S. Skmer: assembly-free and alignment-free sample identification using genome skims. Genome Biol. 2019;20:34.

53. Bernard G, Chan CX, Ragan MA. Alignment-free microbial phylogenomics under scenarios of sequence divergence, genome rearrangement and lateral genetic transfer. Sci Rep. 2016;6:28970.

54. Skippington E, Ragan MA. Within-species lateral genetic transfer and the evolution of transcriptional regulation in Escherichia coli and Shigella. BMC Genomics. 2011;12:532.

55. Beiko RG, Harlow TJ, Ragan MA. Highways of gene sharing in prokaryotes. Proc Natl Acad Sci U S A. 2005;102:14332–7.

56. Darling AE, Miklós I, Ragan MA. Dynamics of genome rearrangement in bacterial populations. PLoS Genet. 2008;4:e1000128.

57. Thankachan SV, Chockalingam SP, Liu Y, Krishnan A, Aluru S. A greedy alignment-free distance estimator for phylogenetic inference. BMC Bioinformatics. 2017;18:238.

58. Doolittle WF, Bapteste E. Pattern pluralism and the Tree of Life hypothesis. Proc Natl Acad Sci U S A. 2007;104:2043–9.

59. Dagan T, Martin W. Getting a better picture of microbial evolution en route to a network of genomes. Philos Trans R Soc Lond B Biol Sci. 2009;364:2187–96.

60. Bernard G, Greenfield P, Ragan MA, Chan CX. k-mer Similarity, Networks of Microbial Genomes, and Taxonomic Rank. mSystems. 2018;3:e00257–18.

61. Bernard G, Ragan MA, Chan CX. Recapitulating phylogenies using -mers: from trees to networks. F1000Res. 2016;5:2789.

62. Earl D, Bradnam K, St John J, Darling A, Lin D, Fass J, et al. Assemblathon 1: a competitive assessment of de novo short read assembly methods. Genome Res. 2011;21:2224–41.

63. Bradnam KR, Fass JN, Alexandrov A, Baranay P, Bechner M, Birol I, et al. Assemblathon 2: evaluating de novo methods of genome assembly in three vertebrate species. Gigascience. 2013;2:10.

64. Baruzzo G, Hayer KE, Kim EJ, Di Camillo B, FitzGerald GA, Grant GR. Simulation-based comprehensive benchmarking of RNA-seq aligners. Nat Methods. 2017;14:135–9.

65. Sczyrba A, Hofmann P, Belmann P, Koslicki D, Janssen S, Dröge J, et al. Critical Assessment of Metagenome Interpretation-a benchmark of metagenomics software. Nat Methods. 2017;14:1063–71.

66. Chandonia J-M, Hon G, Walker NS, Lo Conte L, Koehl P, Levitt M, et al. The ASTRAL Compendium in 2004. Nucleic Acids Res. 2004;32:D189–92.

67. Rice P, Longden I, Bleasby A. EMBOSS: the European Molecular Biology Open Software Suite. Trends Genet. 2000;16:276–7.

68. Tran NH, Chen X. Comparison of next-generation sequencing samples using compression-based distances and its application to phylogenetic reconstruction. BMC Res Notes. 2014;7:320.

69. Hatje K, Kollmar M. A phylogenetic analysis of the brassicales clade based on an alignment-free sequence comparison method. Front Plant Sci. 2012;3:192.

70. Huang W, Li L, Myers JR, Marth GT. ART: a next-generation sequencing read simulator. Bioinformatics. 2012;28:593–4.

71. Beiko RG, Charlebois RL. A simulation test bed for hypotheses of genome evolution. Bioinformatics. 2007;23:825–31.

72. Fan H, Ives AR, Surget-Groba Y, Cannon CH. An assembly and alignment-free method of phylogeny reconstruction from next-generation sequencing data. BMC Genomics. 2015;16:522.

73. Comin M, Antonello M. On the comparison of regulatory sequences with multiple resolution Entropic Profiles. BMC Bioinformatics. 2016;17:130.

74. Fernandes F, Freitas AT, Almeida JS, Vinga S. Entropic Profiler – detection of conservation in genomes using information theory. BMC Res Notes. 2009;2:72.

75. Comin M, Antonello M. Fast Entropic Profiler: An Information Theoretic Approach for the Discovery of Patterns in Genomes. IEEE/ACM Trans Comput Biol Bioinform. 2014;11:500–9.

76. Lau AK, Leimeister C-A, Morgenstern B. Read-SpaM: assembly-free and alignment-free comparison of bacterial genomes with low sequencing coverage. bioRxiv [Internet]. 2019; Available from: https://doi.org/10.1101/550632

77. Gardner SN, Slezak T, Hall BG. kSNP3.0: SNP detection and phylogenetic analysis of genomes without genome alignment or reference genome. Bioinformatics. 2015;31:2877–8.

78. Murray KD, Webers C, Ong CS, Borevitz J, Warthmann N. kWIP: The k-mer weighted inner product, a de novo estimator of genetic similarity. PLoS Comput Biol. 2017;13:e1005727.

79. Filatov G, Bauwens B, Kertész-Farkas A. LZW-Kernel: fast kernel utilizing variable length code blocks from LZW compressors for protein sequence classification. Bioinformatics. 2018;34:3281–8.

80. Stamatakis A. RAxML version 8: a tool for phylogenetic analysis and post-analysis of large phylogenies. Bioinformatics. 2014;30:1312–3.

81. Snir S, Rao S. Quartet MaxCut: a fast algorithm for amalgamating quartet trees. Mol Phylogenet Evol. 2012;62:1–8.

82. Kolekar P, Kale M, Kulkarni-Kale U. Alignment-free distance measure based on return time distribution for sequence analysis: applications to clustering, molecular phylogeny and subtyping. Mol Phylogenet Evol. 2012;65:510–22.

83. Röhling S, Morgenstern B. The number of spaced-word matches between two DNA sequences as a function of the underlying pattern weight [Internet]. bioRxiv. 2019 [cited 2019 Mar 26]. p. 527515. Available from: https://www.biorxiv.org/content/10.1101/527515v1.abstract

84. Leimeister C-A, Boden M, Horwege S, Lindner S, Morgenstern B. Fast alignment-free sequence comparison using spaced-word frequencies. Bioinformatics. 2014;30:1991–9.

85. Horwege S, Lindner S, Boden M, Hatje K, Kollmar M, Leimeister C-A, et al. Spaced words and kmacs: fast alignment-free sequence comparison based on inexact word matches. Nucleic Acids Res. 2014;42:W7–11.

86. Morgenstern B, Zhu B, Horwege S, Leimeister CA. Estimating evolutionary distances between genomic sequences from spaced-word matches. Algorithms Mol Biol. 2015;10:5.

87. Ma B, Tromp J, Li M. PatternHunter: faster and more sensitive homology search. Bioinformatics. 2002;18:440–5.

88. Li M, Ma B, Kisman D, Tromp J. Patternhunter II: highly sensitive and fast homology search. J Bioinform Comput Biol. 2004;02:417–39.

89. Lin J. Divergence measures based on the Shannon entropy [Internet]. IEEE Transactions on Information Theory. 1991. p. 145–51. Available from: http://dx.doi.org/10.1109/18.61115

90. Comin M, Verzotto D. Alignment-free phylogeny of whole genomes using underlying subwords. Algorithms Mol Biol. 2012;7:34.

91. Pedregosa F, Varoquaux G, Gramfort A, Michel V, Thirion B, Grisel O, et al. Scikit-learn: Machine Learning in Python. J Mach Learn Res. 2011;12:2825–30.

92. Huerta-Cepas J, Serra F, Bork P. ETE 3: Reconstruction, Analysis, and Visualization of Phylogenomic Data. Mol Biol Evol. 2016;33:1635–8.

93. Brenner SE, Koehl P, Levitt M. The ASTRAL compendium for protein structure and sequence analysis. Nucleic Acids Res. 2000;28:254–6.

