## Additional file 2 for "Benchmarking of alignment-free sequence comparison methods"

### Additional file 1

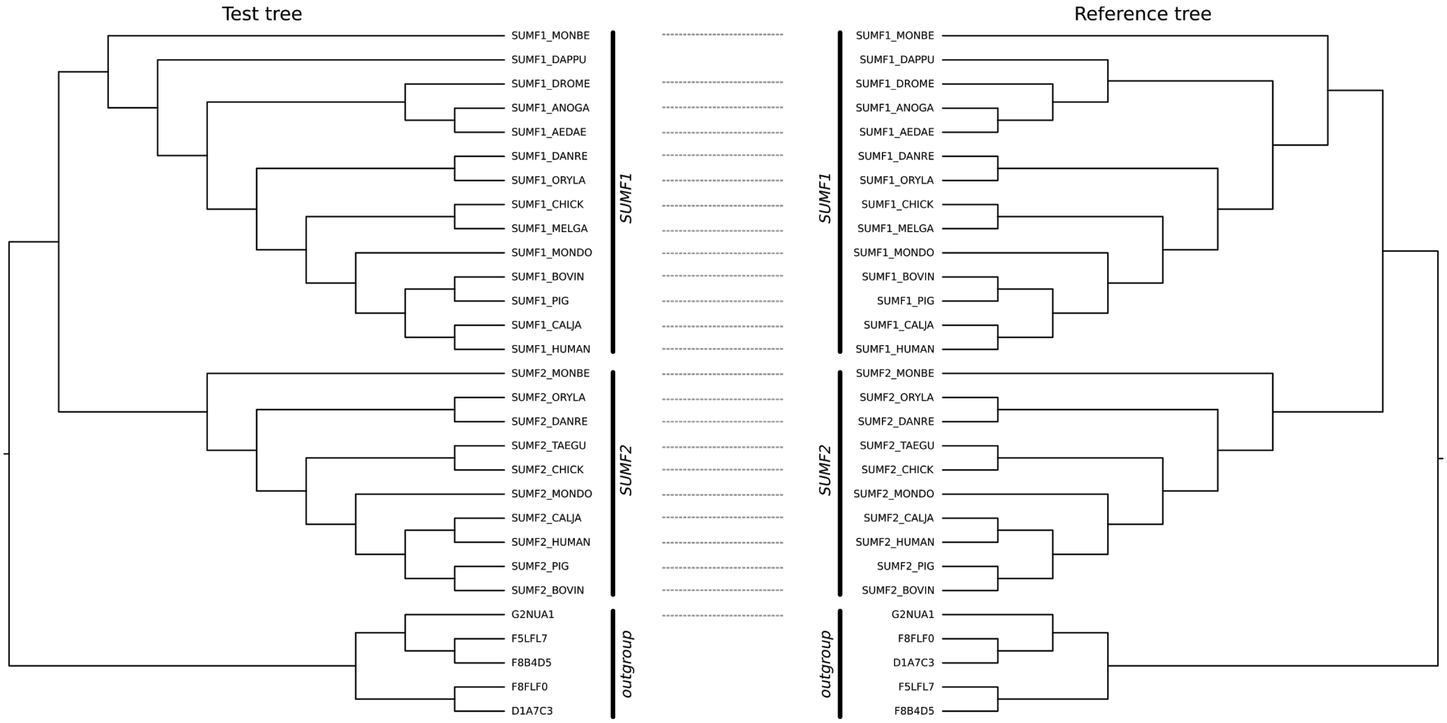


**Figure S1.** Comparison (tanglement) of test tree and reference tree of sulfatase modifying factor (SUMF) gene family in Eukaryotes. The test tree is inferred by 4 alignment-free measures in AFKS program (mismatch, markov, rre_k_r, kl_conditional). Reference phylogenetic tree was taken from SwissTree.

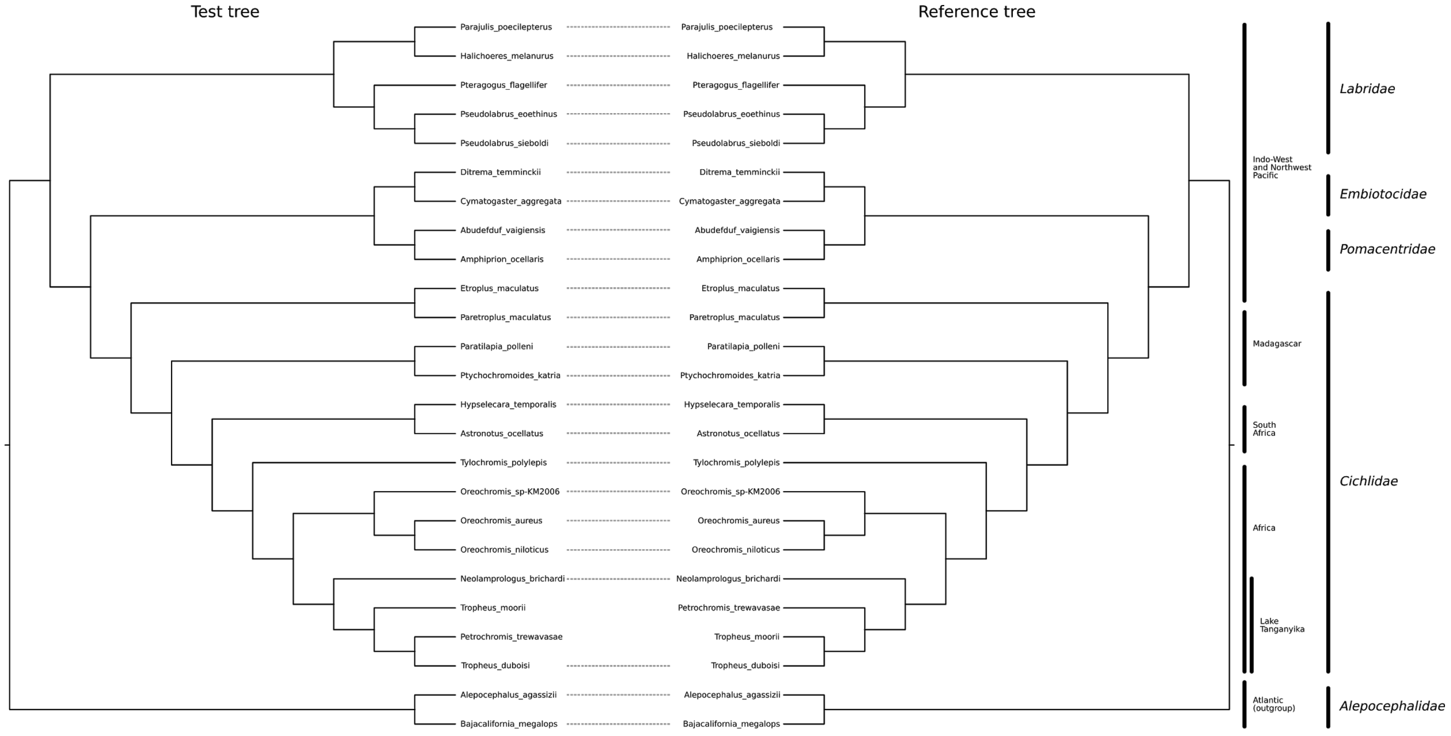


**Figure S2.** Comparison (tanglement) of test tree and reference tree of complete mitochondrial genomes from 25 labroid fishes. The test tree is inferred by 9 alignment-free programs (AFKS, alfpy, CAFE, FSWM, jD2Stat, kmacs, msh, RTD-Phylogeny and spaced). Reference phylogenetic tree was taken from [1].

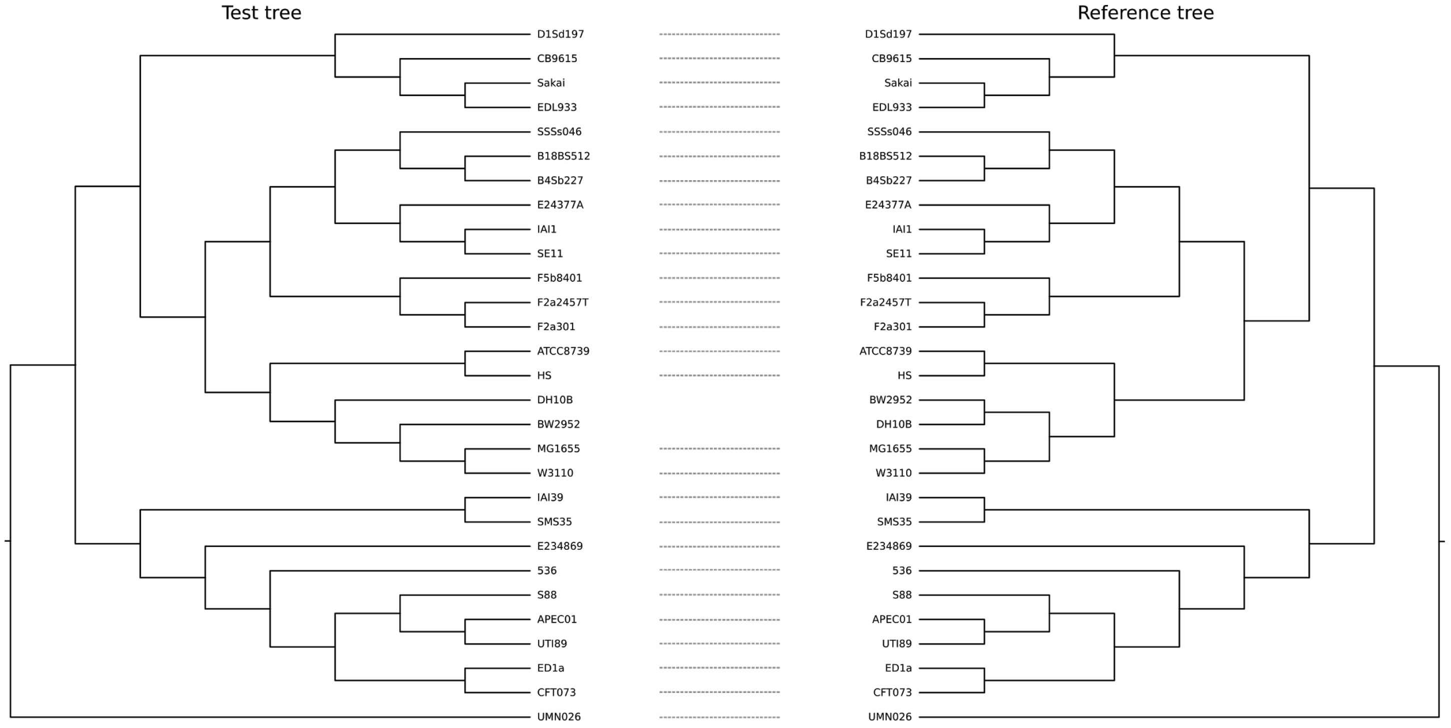


**Figure S3.** Comparison (tanglement) of test and reference cladograms of complete genomes from 29 *E. coli* / *Shigella* species. Test tree was inferred by Phylonium.

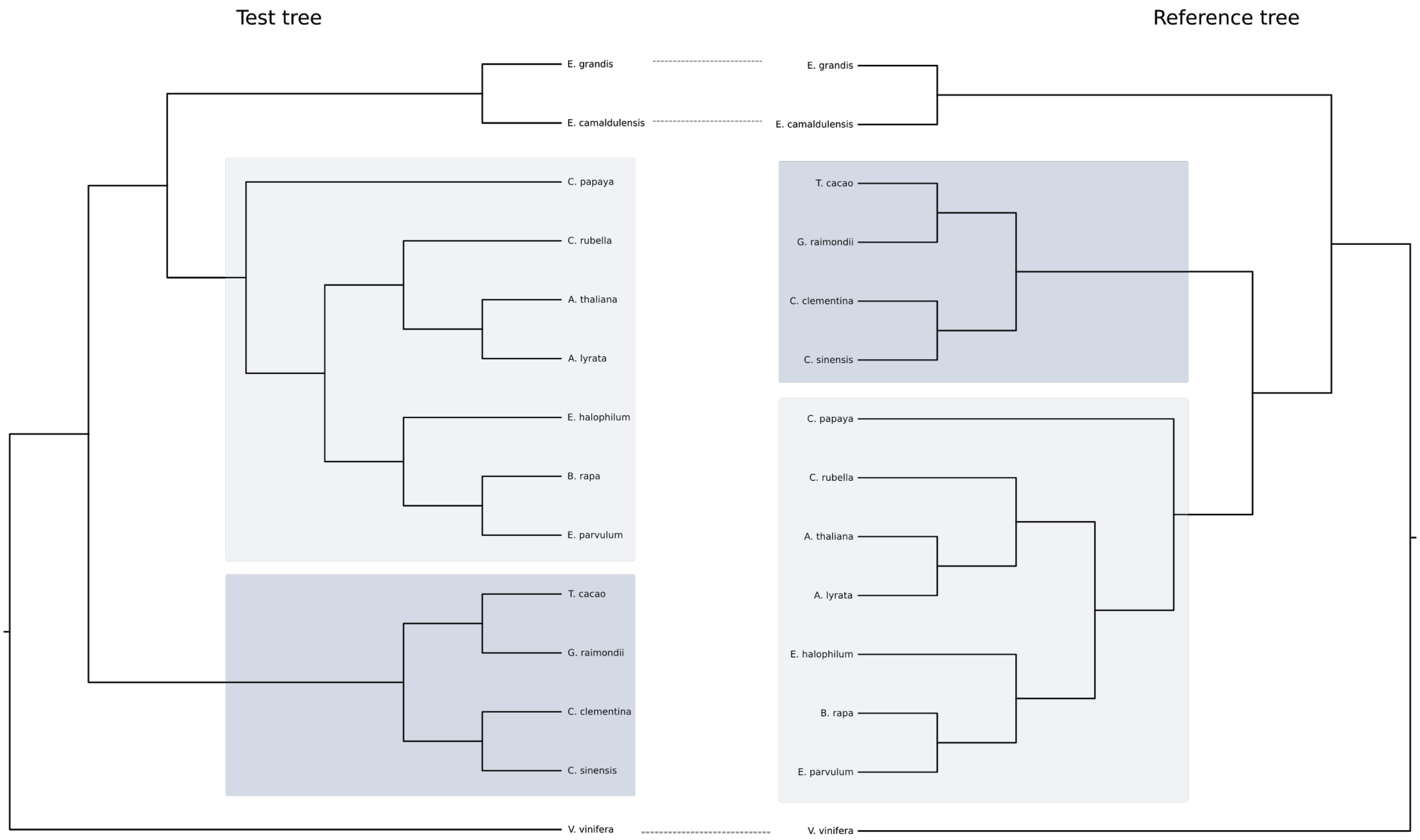


**Figure S4.** Comparison (tanglement) of test and reference cladograms of complete genomes from 14 plant species. Test tree was inferred by Co-Phylog.

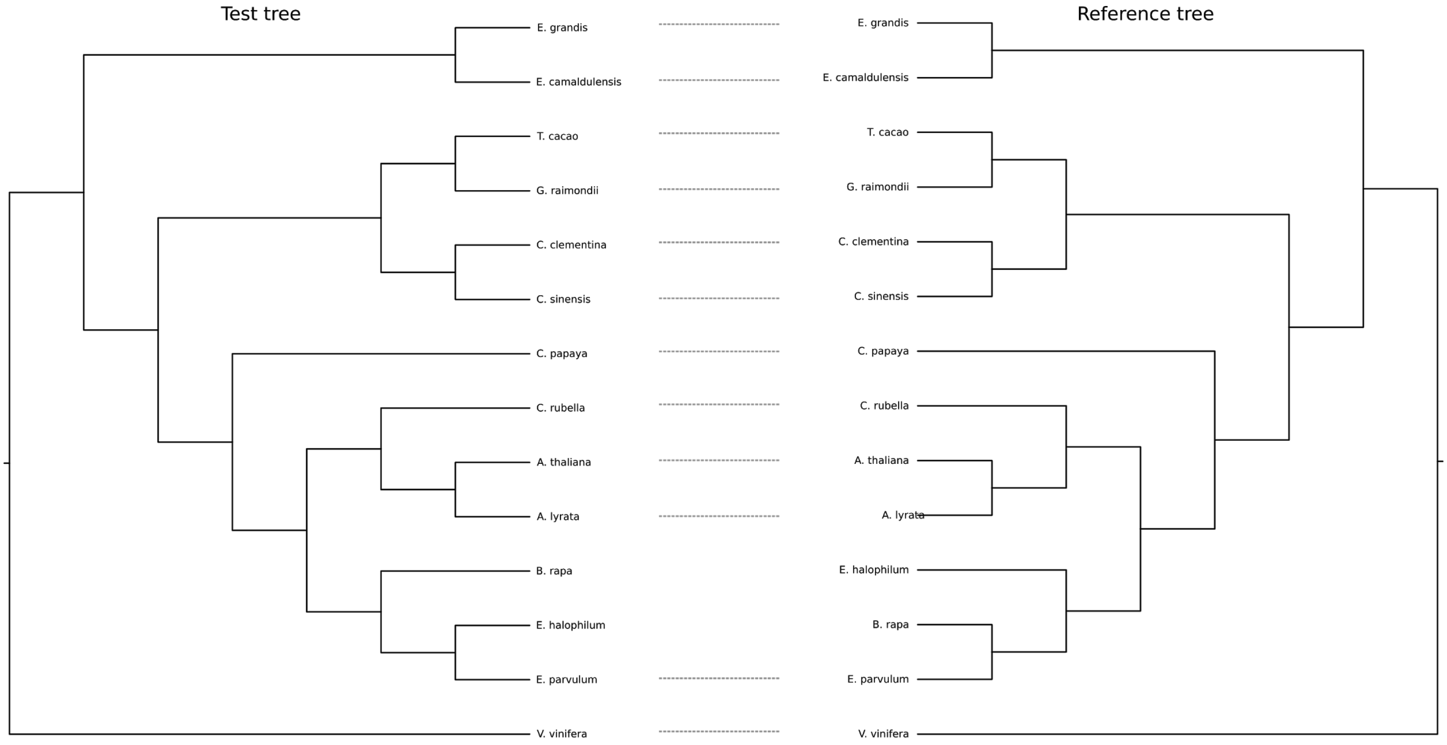


**Figure S5.** Comparison (tanglement) of test and reference cladograms of complete genomes from 14 plant species. Test tree was inferred by mash.

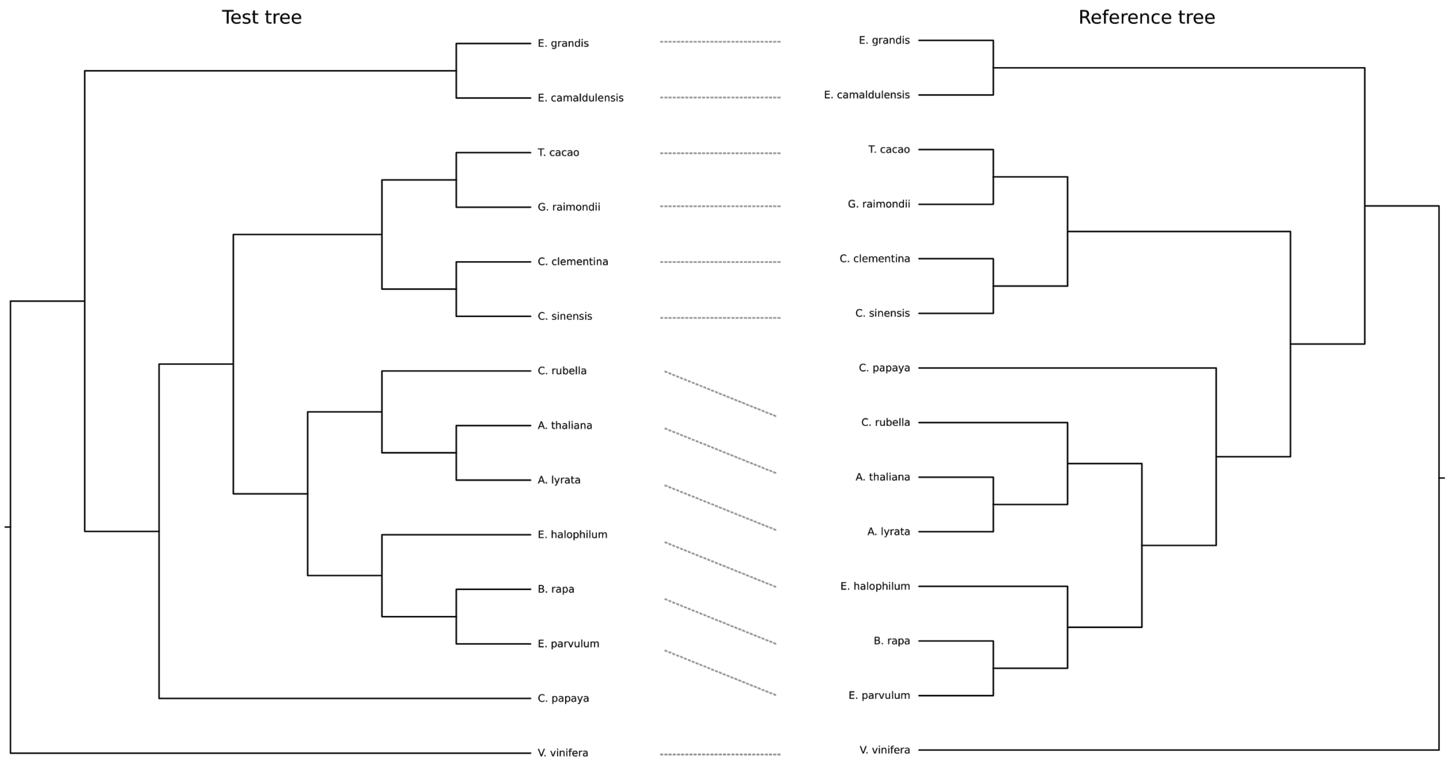


**Figure S6.** Comparison (tanglement) of test and reference cladograms of complete genomes from 14 plant species. Test tree was inferred by Multi-SpaM.

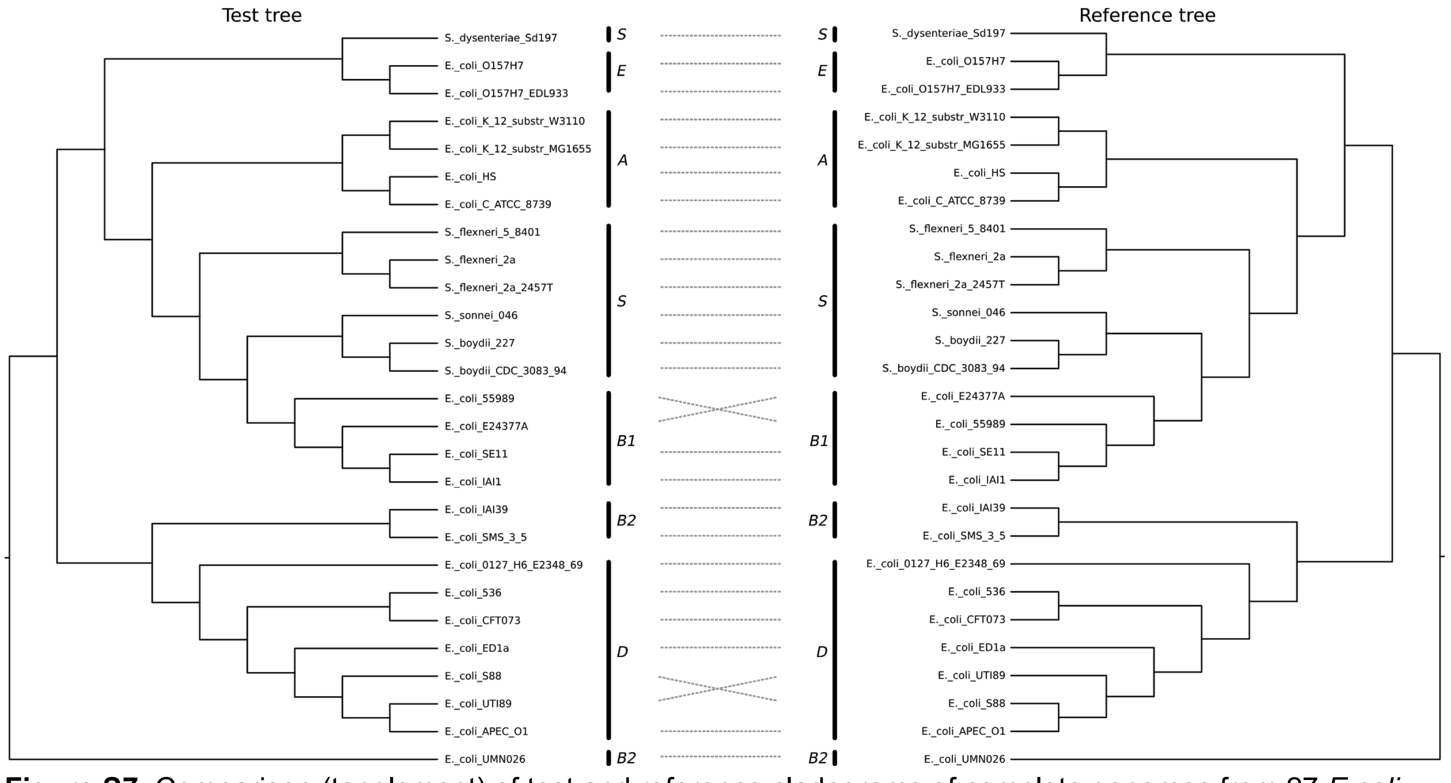


**Figure S7.** Comparison (tanglement) of test and reference cladograms of complete genomes from 27 *E.coli* and *Shigella* species. Test tree was inferred by andi and co-phylog. Reference phylogenetic tree was constructed in [2–4] from 5282 Bayesian protein trees. *E. coli* reference groups and Shigella (S) are indicated.

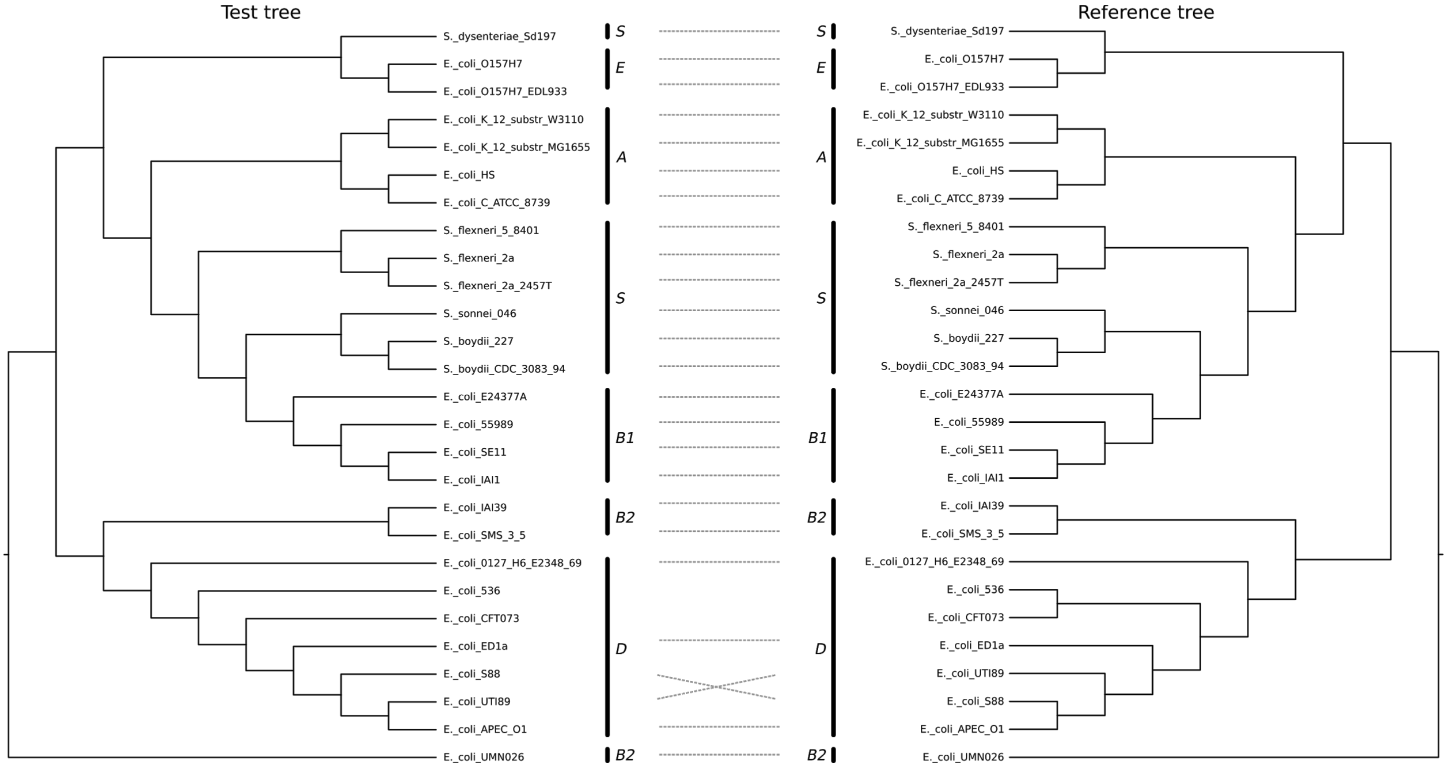


**Figure S8.** Comparison (tanglement) of test and reference cladograms of complete genomes from 27 *E.coli* and *Shigella* species. Test tree was inferred by phylonium. Reference phylogenetic tree was constructed in [2–4] from 5282 Bayesian protein trees. *E. coli* reference groups and Shigella (S) are indicated.

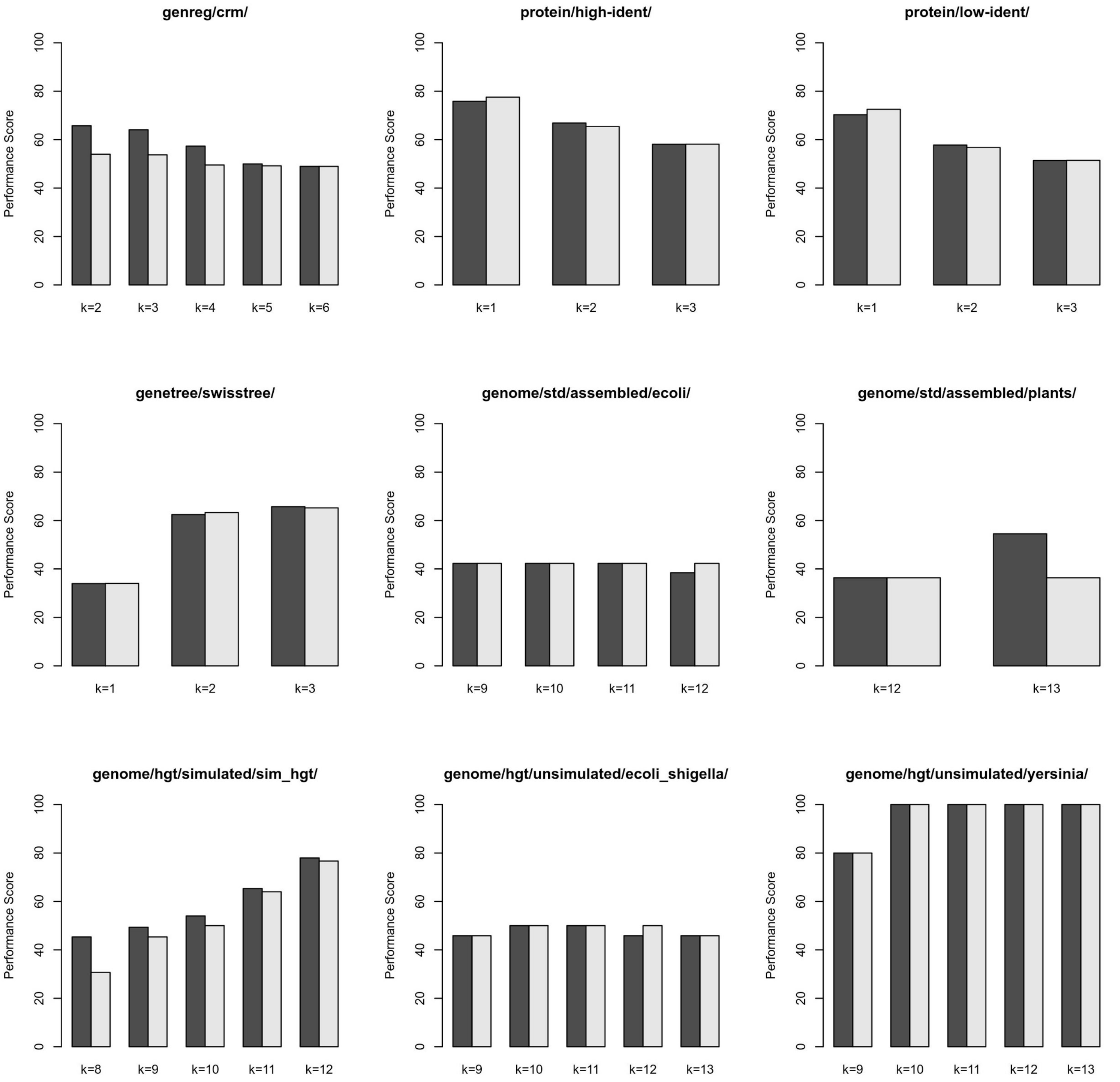


**Figure S9.** Performance scores obtained by alfpy--canberra (dark grey) and AFKS-canberra (light grey) run on common word lengths across datasets.

#### References

1. Fischer C, Koblmüller S, Gülly C, Schlötterer C, Sturmbauer C, Thallinger GG. Complete mitochondrial DNA sequences of the threadfin cichlid (Petrochromis trewavasae) and the blunthead cichlid (Tropheus moorii) and patterns of mitochondrial genome evolution in cichlid fishes. PLoS One. 2013;8:e67048.

2. Skippington E, Ragan MA. Within-species lateral genetic transfer and the evolution of transcriptional regulation in Escherichia coli and Shigella. BMC Genomics. 2011;12:532.

3. Beiko RG, Harlow TJ, Ragan MA. Highways of gene sharing in prokaryotes. Proc Natl Acad Sci U S A. 2005;102:14332–7.

4. Bernard G, Chan CX, Ragan MA. Alignment-free microbial phylogenomics under scenarios of sequence divergence, genome rearrangement and lateral genetic transfer. Sci Rep. 2016;6:28970.
